# Tumor-responsive AAV arms cancer cells to enhance CAR-T cell therapy in solid tumors

**DOI:** 10.64898/2026.11.16.766910

**Authors:** Ziyan Kong, Yile Wang, Yunqi Zhao, Aiwei Yan, Xuhui Wen, Xingyi Xie, Lu Wang, Zhimin Fan, Yongqian Shu, Jinke Wang

## Abstract

Chimeric antigen receptor (CAR)-T-cell therapy remains limited in solid tumors by the scarcity of suitable target antigens and an immunosuppressive tumor microenvironment. Here, we developed a tumor-responsive recombinant adeno-associated virus (rAAV) strategy that harnesses aberrant NF-κB activity to establish a shared synthetic CAR target across tumor types. Using an NF-κB-responsive decoy minimal promoter (DMP), rAAV-SA drove tumor-preferential surface expression of a synthetic antigen (SA), enabling antigen-dependent CAR-T-cell activation, degranulation, and cytotoxicity. Systemic rAAV-SA delivery combined with CAR-T cells produced antitumor activity in triple-negative breast cancer (TNBC) and colorectal cancer xenograft models. We subsequently engineered rAAV-ARMS (antigen redirection and multifunctional secreted effectors), integrating SA expression with tumor-local production of an IL-15/IL-15Rα sushi-domain fusion, CXCL10, and IFN-γ. Compared with antigen redirection alone, rAAV-ARMS enhanced CAR-T-cell-mediated killing in three-dimensional tumor spheroids and, in a TNBC xenograft model, increased intratumoral T-cell accumulation and improved the efficacy of intravenously administered CAR-T cells, without overt systemic toxicity under the conditions examined. These findings establish a tumor-programmed platform for target-redirected universal CAR-T-cell therapy in solid tumors.

## Introduction

Chimeric antigen receptor (CAR)-T-cell therapy has achieved remarkable success in hematological malignancies. However, its efficacy against solid tumors remains limited by the paucity of tumor surface antigens that can be safely and effectively targeted.^1^ To circumvent this antigen bottleneck, recent studies have attempted to install de novo CAR targets in tumors using oncolytic viruses,^2^ engineered bacterial systems,^3^ membrane-inserting ligands,^4^ fusogenic antigen-loaded nanoparticles,^5^ and lipid nanoparticles.^6^ Despite these advances, several challenges persist, including biosafety concerns associated with replication-competent viral and live bacterial vectors,^7,8^ the risk of off-tumor synthetic-antigen expression,^4^ and the maintenance of sufficient antigen expression over time.^6^

Beyond antigen availability, the immunosuppressive tumor microenvironment (TME) represents another major barrier to CAR-T-cell therapy by impairing T-cell infiltration, expansion, and effector function in solid tumors,^9,10^ thereby necessitating complementary strategies to improve therapeutic efficacy. Cytokine-based immunotherapy has emerged as a promising approach to overcome these microenvironmental barriers.^11,12^ Among cytokines, interleukin-15 (IL-15) is particularly attractive because it promotes CD8⁺ T-cell expansion while providing substantially less support for regulatory T-cell (Treg) expansion than IL-2, thereby favoring antitumor T-cell responses.^11^ Moreover, IL-15/IL-15Rα-based superagonists, such as N-803, are designed to mimic physiological IL-15 trans-presentation and exhibit enhanced stability and biological activity.^13,14^ In parallel, the CXCL10–CXCR3 axis provides a major chemotactic signal for activated T-cell trafficking into tumors, whereas IFN-γ can upregulate ICAM-1 on tumor cells and strengthen LFA-1-dependent T-cell–tumor cell adhesion, thereby facilitating CAR-T-cell cytotoxicity.^15,16^ However, systemic cytokine delivery is often constrained by unfavorable pharmacokinetics, widespread peripheral exposure, and treatment-related inflammatory toxicities.^17–19^ These limitations motivate the development of strategies that concentrate complementary immune signals within tumors to support CAR-T-cell function while minimizing systemic exposure.

Adeno-associated virus (AAV) has emerged as a leading gene-delivery platform owing to its favorable safety profile, established clinical utility, and capacity for sustained transgene expression.^20^ However, AAV-mediated delivery alone does not ensure tumor-selective expression, making transcriptional control an important complementary strategy.^21^ Aberrant NF-κB activation, a common feature of many malignancies, provides an endogenous regulatory signal that can be harnessed for tumor-responsive gene expression.^22^ We harnessed this feature to develop an NF-κB-responsive decoy minimal promoter (DMP), consisting of tandem κB-binding elements upstream of a minimal promoter, to preferentially drive therapeutic gene expression in tumor cells.^23–29^ Our previous studies demonstrated the applicability of this regulatory system to diverse molecular modalities, including microRNAs,^25,26^ CRISPR-based effectors,^24^ and proteins,^28,29^ in multiple cellular and animal models.

Building on this platform, we developed a tumor-programmed strategy for universal CAR-T-cell targeting by introducing a shared synthetic antigen (SA) across different solid tumors. We first constructed rAAV-SA, a recombinant AAV vector designed to drive DMP-controlled surface expression of SA, thereby establishing a common target for CAR-T-cell recognition. To further address the immunosuppressive tumor microenvironment, we developed rAAV-ARMS (antigen redirection and multifunctional secreted effectors), which integrates the SA module with DMP-regulated expression of an IL-15/IL-15Rα sushi-domain fusion, CXCL10, and IFN-γ. This tumor-responsive platform combines programmable antigen redirection with multifunctional immune modulation, offering an integrated strategy to overcome both target scarcity and microenvironmental barriers to CAR-T-cell therapy in solid tumors.

## Results

### Engineering tumor-preferential cell-surface display of a synthetic antigen

We first constructed an adeno-associated virus (AAV) plasmid, designated pAAV-DMP-SA, to drive preferential cell-surface expression of a synthetic antigen (SA) in tumor cells (Figure 1A). SA was placed under the control of an NF-κB-responsive decoy minimal promoter (DMP), comprising an NF-κB decoy sequence with five tandem high-affinity NF-κB-binding sites fused to a minimal promoter. In tumor cells with activated NF-κB signaling, NF-κB binds to these sites and activates the adjacent minimal promoter, thereby driving downstream antigen expression. Epidermal growth factor receptor variant III (EGFRvIII) is a tumor-specific deletion mutant of EGFR generated by an in-frame deletion of exons 2–7 and is absent from normal tissues.^30^ A peptide derived from the unique junctional epitope of EGFRvIII, referred to here as PEPvIII, was selected as the model synthetic antigen because it can be recognized by the corresponding CAR-T cells. To enable cell-surface localization, PEPvIII was linked to a platelet-derived growth factor receptor (PDGFR) transmembrane domain, with a Myc tag and a flexible linker incorporated into the construct. We next examined the expression of RELA, which encodes the p65 subunit of NF-κB, in a panel of tumor cell lines (MDA-MB-468, PANC-1, HCT116, MDA-MB-231, and MCF-7) and non-tumor cell lines (MCF-12A, MRC-5, GES-1, HL7702, and NIH-3T3). RELA mRNA levels were significantly higher in the tumor cell lines than in the non-tumor cell lines (Figure 1B). Tumor and non-tumor cells were then transfected with the pAAV-DMP-SA plasmid or the empty control vector pAAV-MCS, and cell-surface PEPvIII expression was evaluated by flow cytometry 72 h after transfection. Transfection with pAAV-DMP-SA significantly increased the proportion of PEPvIII-positive cells in MDA-MB-231, MCF-7, PANC-1, and HCT116 cells, while PEPvIII-positive populations remained near background levels in MCF-12A, MRC-5, GES-1, and HL7702 cells (Figure 1C). Together, these results indicate that pAAV-DMP-SA preferentially introduces a synthetic antigen onto the surface of multiple tumor cell lines, thereby generating a de novo target for CAR-T-cell recognition—a strategy of particular relevance to triple-negative breast cancer, in which suitable tumor-specific surface targets remain scarce.

**Figure 1.**
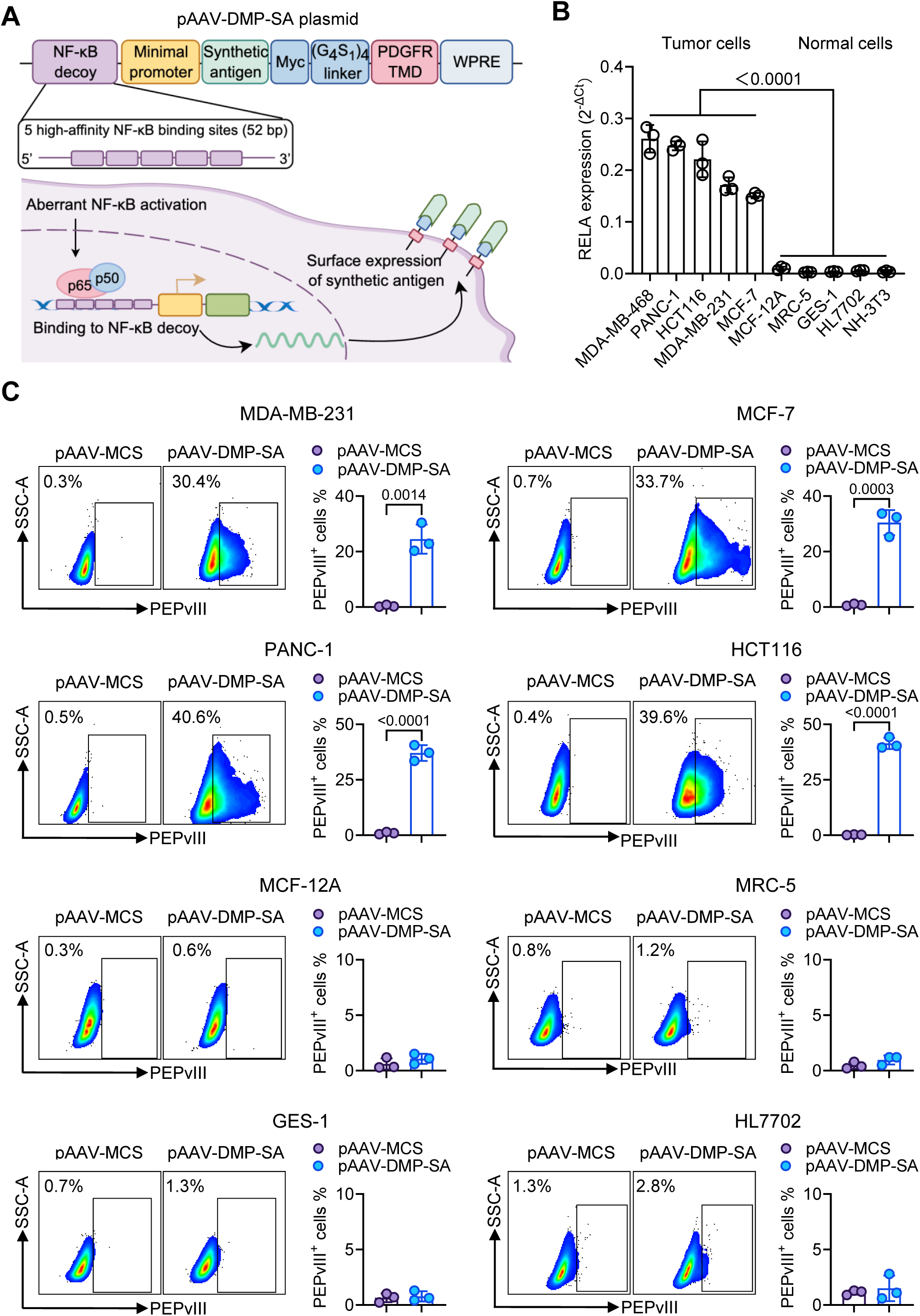
DMP-driven tumor-preferential cell-surface expression of a synthetic antigen. (A) Schematic of the pAAV-DMP-SA expression strategy. The synthetic antigen was placed under the control of an NF-κB-responsive decoy minimal promoter (DMP) containing five tandem high-affinity NF-κB-binding sites, with a PDGFR transmembrane domain for cell-surface localization and WPRE positioned downstream to enhance expression. In tumor cells with activated NF-κB signaling, NF-κB binds to the DMP and drives synthetic antigen expression on the cell surface. (B) RELA (p65) mRNA expression in tumor (MDA-MB-468, PANC-1, HCT116, MDA-MB-231, and MCF-7) and non-tumor (MCF-12A, MRC-5, GES-1, HL7702, and NIH-3T3) cell lines. (C) Flow-cytometry analysis of cell-surface PEPvIII expression in tumor (MDA-MB-231, MCF-7, PANC-1, and HCT116) and non-tumor (MCF-12A, MRC-5, GES-1, and HL7702) cell lines 72 h after transfection with pAAV-DMP-SA or pAAV-MCS. Representative flow cytometry plots are shown on the left, with corresponding quantification of PEPvIII-positive cells on the right. Data are presented as mean ± SD from three independent experiments. Statistical significance was assessed using unpaired two-tailed Student’s t-tests. P values are indicated in the figure.

### Synthetic antigen redirection enables CAR-T-cell activation and tumor-cell killing

We next investigated whether DMP-driven synthetic antigen expression could render tumor cells targetable by anti-EGFRvIII CAR-T cells. As illustrated in Figure 2A, elevated NF-κB activity drives surface display of the synthetic antigen, thereby converting an intrinsic tumor-associated signaling state into a CAR-binding target on the tumor-cell surface. Antigen–CAR engagement subsequently initiates CAR-T-cell activation and effector responses, culminating in tumor-cell killing. To evaluate this strategy, we constructed an anti-EGFRvIII CAR comprising a CD8α signal peptide, an EGFRvIII-specific single-chain variable fragment (scFv), a CD8 hinge, a CD28 transmembrane domain, and 4-1BB/CD3ζ intracellular signaling domains, with EGFP co-expressed through a P2A sequence (Figure 2B). T cells isolated from human peripheral blood showed approximately 99% CD3⁺ purity before CAR-T-cell generation (Supplementary Figure 1) and were subsequently transduced with CAR-encoding lentivirus. EGFP served as a reporter of transduction efficiency, whereas CAR surface expression was assessed using a fluorescent PEPvIII probe generated by pre-complexing biotinylated PEPvIII with PE-conjugated streptavidin. Approximately 53% of T cells bound the PEPvIII probe, closely matching the EGFP-positive fraction (∼57%), confirming surface expression of antigen-binding anti-EGFRvIII CARs (Figure 2B).

**Figure 2.**
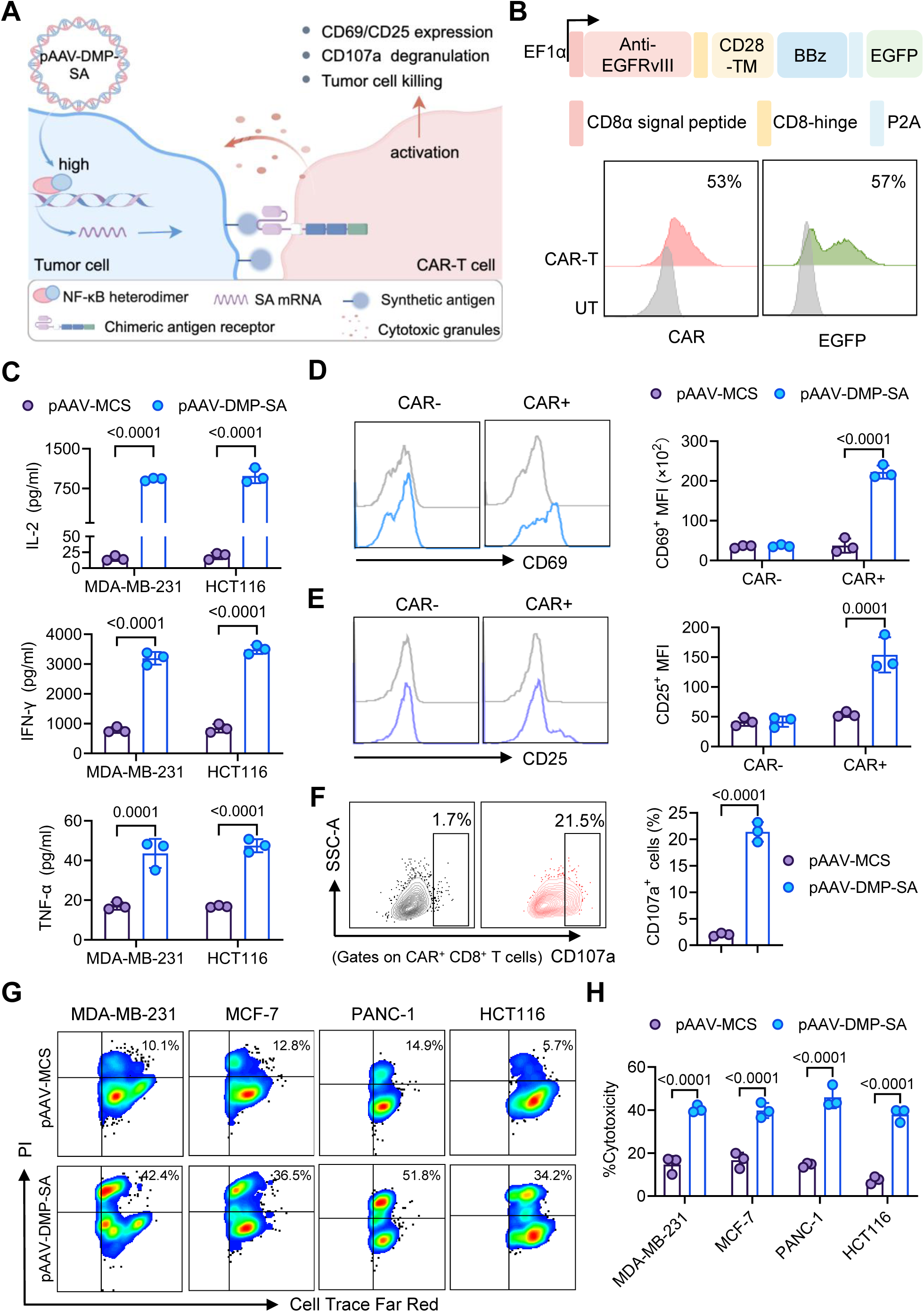
Synthetic antigen expression redirects CAR-T-cell responses and promotes tumor-cell killing. (A) Schematic of the synthetic antigen-mediated CAR-T-cell redirection strategy. Elevated NF-κB activity in tumor cells drives DMP-dependent surface expression of the synthetic antigen, enabling CAR engagement and subsequent T-cell activation, effector responses, and tumor-cell killing. (B) Structure of the anti-EGFRvIII CAR and representative flow cytometry analysis of CAR surface expression and EGFP expression in CAR-T cells compared with untransduced T cells (UT). (C) MDA-MB-231 and HCT116 cells transfected with pAAV-MCS or pAAV-DMP-SA for 72 h were cocultured with CAR-T cells at an effector-to-target (E:T) ratio of 5:1 for 18 h. IL-2, IFN-γ, and TNF-α levels in culture supernatants were measured. (D, E) Representative histograms and quantification of CD69 (D) and CD25 (E) expression in CAR⁻ and CAR⁺ T-cell populations following coculture with pAAV-MCS- or pAAV-DMP-SA-transfected MDA-MB-231 cells. (F) Representative flow cytometry analysis and quantification of CD107a⁺ cells within the CAR⁺CD8⁺ T-cell population following coculture with pAAV-MCS- or pAAV-DMP-SA-transfected MDA-MB-231 cells. (G) Representative flow cytometry analysis of CAR-T-cell-mediated tumor-cell killing. CellTrace Far Red-labeled target cells were distinguished from effector T cells, and dead tumor cells were identified as PI⁺ cells within the CellTrace Far Red⁺ population. Upper and lower rows represent cocultures with pAAV-MCS- and pAAV-DMP-SA-transfected tumor cells, respectively. (H) Quantification of CAR-T-cell-mediated cytotoxicity against MDA-MB-231, MCF-7, PANC-1, and HCT116 cells transfected with pAAV-MCS or pAAV-DMP-SA. Data are presented as mean ±SD from three independent experiments. Statistical significance was assessed by two-way ANOVA followed by Šídák’s multiple-comparisons test (C–E, H) or an unpaired two-tailed Student’s *t*-test (F). P values are shown in the figure.

Recognition of the introduced antigen elicited a pronounced CAR-T-cell cytokine response. CAR-T cells cocultured with pAAV-DMP-SA-transfected MDA-MB-231 and HCT116 cells secreted significantly higher levels of IL-2, IFN-γ, and TNF-α than those exposed to the corresponding pAAV-MCS controls (Figure 2C). We further examined CAR-dependent T-cell activation by analyzing CD69 and CD25 separately in CAR⁺ and CAR⁻ populations. Both markers were increased in CAR⁺ T cells exposed to pAAV-DMP-SA-transfected MDA-MB-231 cells, whereas little change was observed in the CAR⁻ population (Figure 2D, E), demonstrating that the activation response was dependent on CAR recognition of the introduced antigen. We next assessed cytotoxic effector function. CD107a surface mobilization was increased in CAR⁺CD8⁺ T cells exposed to pAAV-DMP-SA-transfected targets, indicating enhanced degranulation (Figure 2F and Supplementary Figure 2A). Target-cell killing was then directly evaluated by distinguishing CellTrace Far Red-labeled tumor cells from effector T cells and quantifying PI uptake within the target-cell population (Supplementary Figure 2B). Representative flow cytometry plots showed higher proportions of PI⁺ target cells following pAAV-DMP-SA transfection in MDA-MB-231, MCF-7, PANC-1, and HCT116 cells (Figure 2G). Quantitative analysis further confirmed significantly greater CAR-T-cell-mediated cytotoxicity across all four tumor cell lines compared with their respective pAAV-MCS controls (Figure 2H). Together, these findings demonstrate that DMP-driven PEPvIII surface display converts NF-κB activity into a targetable surface signal, enabling antigen-dependent CAR-T-cell cytokine secretion, activation, degranulation, and direct tumor-cell killing across multiple tumor models, including triple-negative breast cancer.

### rAAV-mediated antigen redirection enables CAR-T-cell antitumor activity in vivo

To translate the DMP-SA strategy to in vivo delivery, we next packaged the DMP-controlled expression cassette into AAV2 vectors. Because subsequent treatment involved CAR-T cells, unintended synthetic-antigen expression in T cells could potentially lead to CAR-T-cell fratricide. To assess transgene expression in these cells, we compared rAAV-DMP-ZsGreen-mediated reporter expression in unstimulated T cells, activated T cells, and MDA-MB-231 cells by flow cytometry. ZsGreen fluorescence remained at background levels in both T-cell populations, whereas MDA-MB-231 cells displayed a significant increase in ZsGreen MFI (Supplementary Figure 3). These findings indicate limited detectable reporter expression in primary T cells under the experimental conditions, supporting further evaluation of this vector system in combination with CAR-T cells.

We next evaluated whether systemic rAAV-SA delivery could establish synthetic antigen expression in tumors in vivo. When tumors reached approximately 100 mm³on day 25, MDA-MB-231 tumor-bearing NCG mice received a single intravenous dose of rAAV-MCS or rAAV-SA (5 × 10¹⁰ vg per mouse), and tumors and major organs were collected 7 days later (Figure 3A). Compared with the rAAV-MCS control, rAAV-SA administration alone did not significantly alter endpoint tumor weight or volume (Supplementary Figure 4A–C). Biodistribution analysis showed that rAAV DNA was detectable across the examined tissues, whereas PEPvIII mRNA was predominantly detected in tumor tissue (Supplementary Figure 4D, E). Following sequential gating for cells, singlets, and viable cells (Supplementary Figure 4F), flow cytometric analysis of dissociated tumors demonstrated increased surface PEPvIII expression after rAAV-SA administration, whereas the rAAV-MCS group remained at background levels (Figure 3B). Consistent with these findings, immunohistochemical staining for the Myc tag incorporated into the PDGFR membrane-anchoring scaffold showed clear membrane-associated staining in tumor tissue (Figure 3C), whereas no appreciable DAB staining was detected in the heart, liver, spleen, lung, or kidney (Supplementary Figure 4G). Together, these results demonstrate that systemic rAAV-SA delivery results in preferential synthetic antigen expression in MDA-MB-231 tumors despite broad rAAV vector biodistribution.

**Figure 3.**
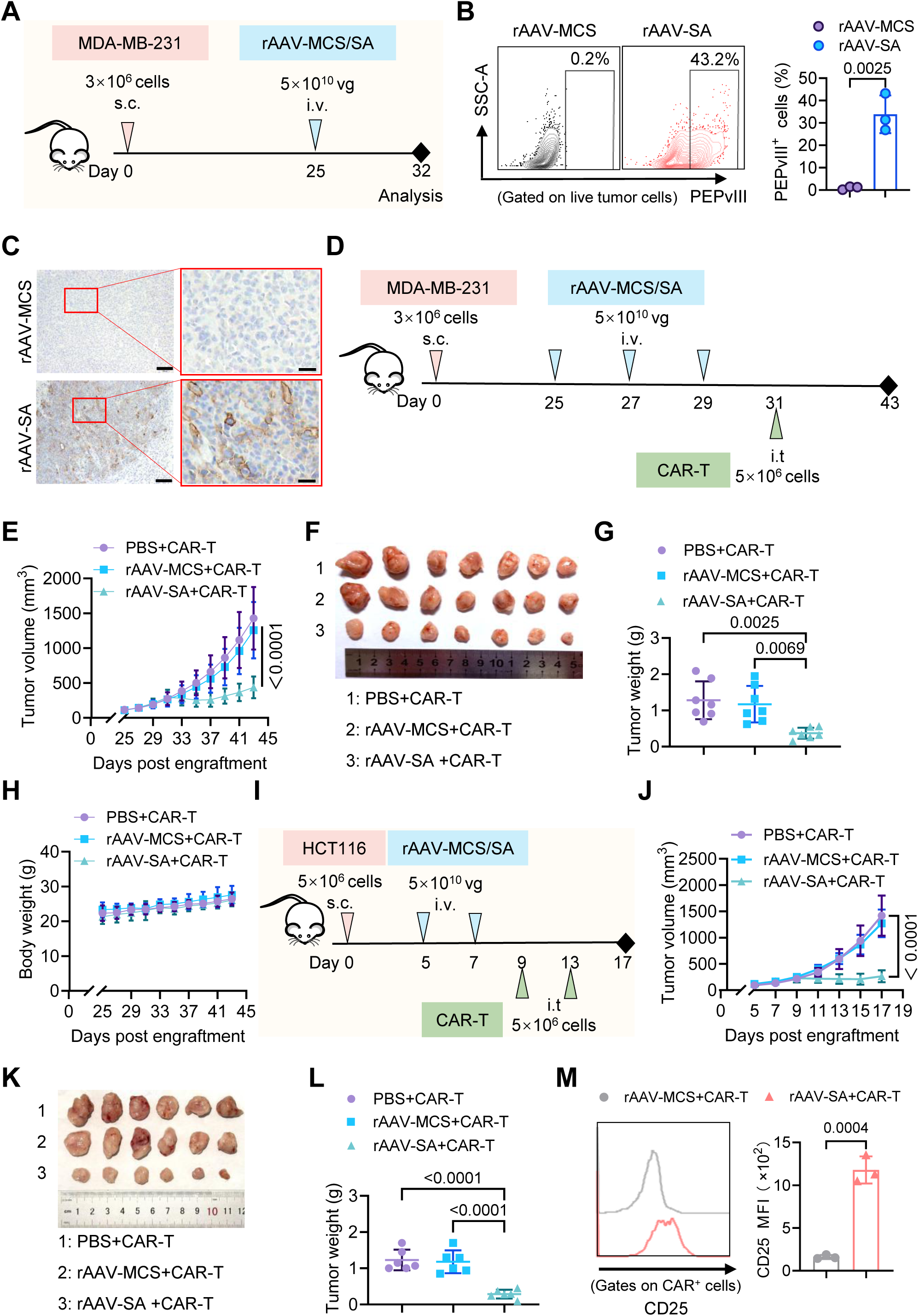
rAAV-delivered synthetic antigen enables CAR-T-cell antitumor activity in vivo. (A–C) Synthetic antigen expression in the MDA-MB-231 xenograft model. When tumors reached approximately 100 mm³, mice received a single intravenous dose of rAAV-MCS or rAAV-SA (5 × 10¹⁰ vg/mouse) on day 25, and tumors were collected one week later. Experimental scheme (A), representative flow cytometry plots and quantification of PEPvIII⁺ cells among live tumor cells (B), and representative Myc-tag immunohistochemistry images (C) are shown (n = 3 mice per group). Scale bars, 100 μm for overview images and 25 μm for magnified images. (D–H) Therapeutic evaluation in the MDA-MB-231 xenograft model. When tumors reached approximately 100 mm³, mice received intravenous injections of PBS, rAAV-MCS, or rAAV-SA (5 × 10¹⁰ vg/mouse) on days 25, 27, and 29, followed by intratumoral CAR-T-cell administration (5 × 10⁶ cells/mouse) on day 31. Treatment scheme (D), tumor growth curves (E), excised tumors (F), endpoint tumor weights (G), and body-weight changes (H) are shown (n = 7 mice per group). (I–L) Therapeutic evaluation in the HCT116 xenograft model. When tumors reached approximately 100 mm³, mice received intravenous injections of PBS, rAAV-MCS, or rAAV-SA (5 × 10¹⁰ vg/mouse) on days 5 and 7, followed by intratumoral CAR-T-cell administration (5 × 10⁶ cells/mouse) on days 9 and 13. Treatment scheme (I), tumor growth curves (J), excised tumors (K), and endpoint tumor weights (L) are shown (n = 6 mice per group). (M) Representative flow cytometry histograms and quantification of CD25 mean fluorescence intensity (MFI) in intratumoral CAR⁺ T cells from a short-term HCT116 study. When tumors reached approximately 100 mm³, mice received intravenous injections of rAAV-MCS or rAAV-SA (5 × 10¹⁰ vg/mouse) on days 5 and 7, followed by intratumoral CAR-T-cell administration (5 × 10⁶ cells/mouse) on day 9, and tumors were collected two days later (n = 3 mice per group). Data are presented as mean ±SD. Statistical significance was assessed using unpaired two-tailed Student’s t-tests for (B) and (M), two-way ANOVA for (E) and (J), and one-way ANOVA followed by Tukey’s multiple-comparisons test for (G) and (L). P values are shown in the figure.

With in vivo antigen expression established, we evaluated the therapeutic efficacy of rAAV-SA in combination with CAR-T cells in the MDA-MB-231 model. When tumors reached approximately 100 mm³, MDA-MB-231 tumor-bearing NCG mice were treated with PBS, rAAV-MCS, or rAAV-SA (5 × 10¹⁰ vg/mouse per rAAV injection) intravenously on days 25, 27, and 29, followed by intratumoral administration of CAR-T cells (5 × 10⁶ cells/mouse) on day 31 (Figure 3D). Tumors continued to progress in mice receiving CAR-T cells following PBS or rAAV-MCS administration, whereas rAAV-SA combined with CAR-T cells significantly inhibited tumor growth (Figure 3E and Supplementary Figure 5A). At the experimental endpoint, tumors from mice treated with rAAV-SA and CAR-T cells were visibly smaller and exhibited significantly lower weights than those from both control groups (Figure 3F, G). Body weight remained stable throughout treatment, with no significant weight loss observed in any group (Figure 3H). In addition, serum levels of ALT, AST, ALP, BUN, creatinine, and uric acid showed no significant differences among treatment groups (Supplementary Figure 5B), and histological examination revealed no obvious pathological abnormalities in the heart, liver, spleen, lung, or kidney (Supplementary Figure 5C). Together, these findings demonstrate that rAAV-SA-mediated synthetic antigen expression enables CAR-T-cell antitumor activity in vivo without evident treatment-related systemic toxicity under the conditions examined.

To determine whether the therapeutic effect extended beyond the TNBC model, we next evaluated rAAV-SA combined with CAR-T cells in HCT116 colorectal cancer xenografts. When tumors reached approximately 100 mm³, HCT116 tumor-bearing NCG mice received intravenous injections of PBS, rAAV-MCS, or rAAV-SA (5 × 10¹⁰ vg/mouse per rAAV injection) on days 5 and 7, followed by intratumoral CAR-T-cell administration (5 × 10⁶ cells/mouse per injection) on days 9 and 13 (Figure 3I). Similar to the MDA-MB-231 model, tumors in mice receiving CAR-T cells following PBS or rAAV-MCS administration continued to progress, whereas rAAV-SA combined with CAR-T cells significantly inhibited HCT116 tumor growth (Figure 3J and Supplementary Figure 6A). Consistent with the growth curves, tumors from mice receiving rAAV-SA combined with CAR-T cells were visibly smaller and exhibited significantly lower endpoint weights than those from both control groups (Figure 3K, L). We further examined vector biodistribution and synthetic antigen expression in this model. rAAV DNA was detected in multiple tissues, whereas PEPvIII mRNA was detected predominantly in tumors (Supplementary Figure 6B, C). Myc immunohistochemistry likewise revealed clear membrane-associated staining in rAAV-SA-treated tumors (Supplementary Figure 6D), with no appreciable staining in the heart, liver, spleen, lung, or kidney (Supplementary Figure 6E). No significant body-weight loss was observed during treatment (Supplementary Figure 7A), and serum biochemical parameters (Supplementary Figure 7B) and histological examination of major organs (Supplementary Figure 7C) revealed no evident treatment-related abnormalities. These findings further support the antitumor efficacy of rAAV-SA combined with CAR-T cells in distinct solid-tumor xenograft models, without overt systemic toxicity under the conditions examined.

We further investigated whether tumor control was accompanied by local T-cell activation in a short-term HCT116 study. When tumors reached approximately 100 mm³, mice received intravenous rAAV-MCS or rAAV-SA (5 × 10¹⁰ vg/mouse per injection) on days 5 and 7, followed by intratumoral administration of CAR-T cells (5 × 10⁶ cells/mouse) on day 9, and tumors were collected two days later for flow cytometric analysis (Supplementary Figure 8A). CD69 expression was significantly increased in intratumoral CD3⁺ T cells from rAAV-SA-treated mice compared with the rAAV-MCS group (Supplementary Figure 8B). Following identification of intratumoral CAR⁺ T cells using the gating strategy shown in Supplementary Figure 8C, CAR⁺ T cells from rAAV-SA-treated tumors exhibited significantly higher CD25 expression (Figure 3M). Together, these results show that rAAV-SA provides tumor cells with a de novo CAR-recognizable target, promoting local T-cell activation and CAR-T-cell antitumor activity in vivo.

### Multifunctional rAAV-ARMS further enhances CAR-T-cell activity in tumor spheroids

Although synthetic antigen expression enabled CAR-T-cell recognition and antitumor activity, the immunosuppressive tumor microenvironment presents additional barriers to sustained T-cell function. We therefore engineered rAAV-ARMS (antigen redirection and multifunctional secreted effectors), encoding PEPvIII, an IL-15/IL-15Rα sushi-domain fusion (IL-15 complex), CXCL10, and IFN-γ (Figure 4A). Having established synthetic antigen-mediated CAR-T-cell activity in two-dimensional cultures, we next evaluated rAAV-ARMS in MDA-MB-231 and HCT116 three-dimensional tumor spheroids established from 1 × 10³cells per well over 7 days and 48 h, respectively, and treated with rAAV-MCS, rAAV-SA, or rAAV-ARMS (1 × 10⁸ vg/well) for 72 h. Following spheroid dissociation into single-cell suspensions, flow-cytometric analysis revealed that both rAAV-SA and rAAV-ARMS induced surface PEPvIII expression, whereas rAAV-MCS remained at background levels (Figure 4B, C), indicating that synthetic antigen display was maintained following incorporation of the additional payloads. ELISA further detected the IL-15 complex, CXCL10, and IFN-γ in supernatants from rAAV-ARMS-treated spheroids, whereas these factors were undetectable in rAAV-MCS-treated controls (Figure 4D).

**Figure 4.**
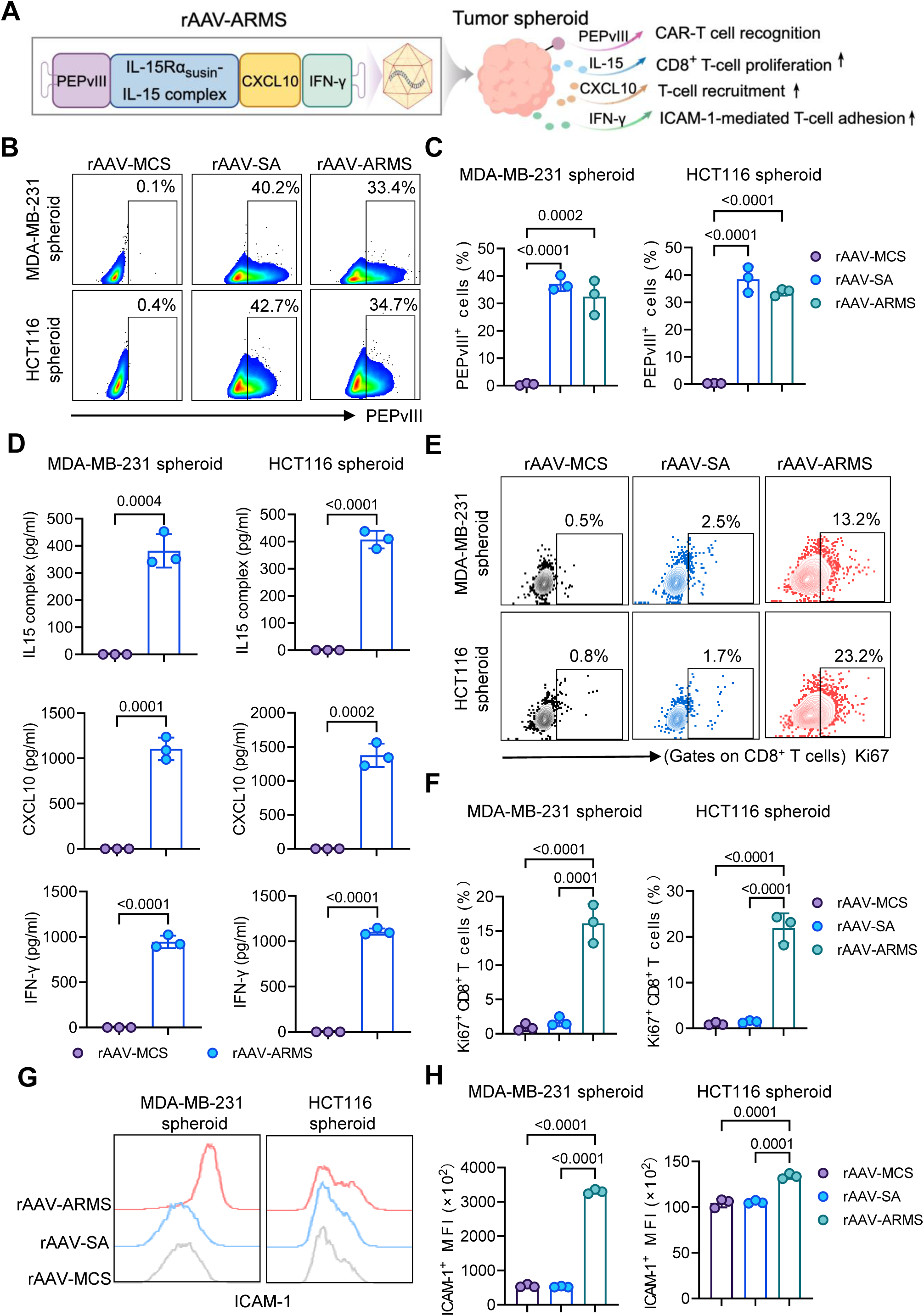
Construction and functional characterization of rAAV-ARMS in tumor spheroids. (A) Schematic of rAAV-ARMS, which encodes PEPvIII, an IL-15/IL-15Rα sushi-domain fusion (IL-15 complex), CXCL10, and IFN-γ to combine synthetic antigen display with additional immune-supportive components. (B) Representative flow cytometry plots of surface PEPvIII expression in MDA-MB-231 and HCT116 spheroid-derived cells following treatment with rAAV-MCS, rAAV-SA, or rAAV-ARMS. (C) Quantification of PEPvIII⁺ cells in the indicated spheroid-derived cell populations. (D) Concentrations of IL-15 complex, CXCL10, and IFN-γ in culture supernatants from rAAV-MCS or rAAV-ARMS-treated spheroids measured by ELISA. (E) Representative flow cytometry plots of Ki67 expression in CD8⁺ primary human T cells cultured for 48 h with conditioned media collected from the indicated spheroids. (F) Quantification of Ki67⁺ cells among CD8⁺ T cells. (G) Representative flow cytometry histograms of surface ICAM-1 expression in MDA-MB-231 and HCT116 spheroid-derived cells following the indicated rAAV treatments. (H) Quantification of ICAM-1 mean fluorescence intensity (MFI). Data are presented as mean ± SD from three independent experiments. Statistical significance was assessed using an unpaired two-tailed Student’s *t*-test for (D) and one-way ANOVA with multiple-comparisons testing for (C), (F), and (H). P values are shown in the figure.

To assess whether factors secreted by rAAV-ARMS-treated spheroids could influence T-cell proliferation, primary human T cells were cultured for 48 h with conditioned media collected from the indicated spheroids. Conditioned media from rAAV-ARMS-treated spheroids significantly increased the proportion of Ki67⁺ cells among CD8⁺ T cells compared with media from rAAV-MCS- or rAAV-SA-treated spheroids (Figure 4E, F), consistent with the known proliferative effect of the IL-15 complex on T cells. To further characterize tumor-cell responses to rAAV-ARMS, rAAV-treated spheroids were dissociated into single-cell suspensions for flow-cytometric analysis of surface ICAM-1 expression. rAAV-ARMS significantly increased ICAM-1 expression on both MDA-MB-231 and HCT116 cells relative to rAAV-MCS and rAAV-SA (Figure 4G, H), consistent with IFN-γ-associated ICAM-1 upregulation. Together, these findings show that rAAV-ARMS preserves synthetic antigen display while enabling secretion of the incorporated effector molecules and promoting CD8⁺ T-cell proliferation and tumor-cell ICAM-1 upregulation.

We next asked whether the cellular changes induced by rAAV-ARMS translated into enhanced CAR-T-cell killing in three-dimensional tumor models. To assess potential direct effects of rAAV treatment on tumor spheroids, MDA-MB-231 and HCT116 spheroids were treated with PBS, rAAV-MCS, rAAV-SA, or rAAV-ARMS in the absence of T cells. No significant differences in spheroid diameter or cell viability were observed after 72 h, as assessed by bright-field imaging, Calcein AM/PI/Hoechst staining, and luciferase-based viability analysis (Supplementary Figure 9A–F), indicating that rAAV treatment alone did not exert detectable direct cytotoxic effects on tumor spheroids under these in vitro conditions.

Following CAR-T-cell coculture at an effector-to-target (E:T) ratio of 10:1 for 48 h, both rAAV-SA and rAAV-ARMS significantly reduced HCT116 spheroid diameter compared with rAAV-MCS, whereas no significant difference was detected between the rAAV-SA and rAAV-ARMS groups (Figure 5A, B). We next evaluated tumor-cell death more directly. Bright-field imaging suggested increased cell accumulation around rAAV-ARMS-treated spheroids following CAR-T-cell coculture (Figure 5C, D). Caspase-3/7 imaging of spheroids prelabeled with CellTracker Blue revealed increased apoptotic signals following rAAV-SA or rAAV-ARMS treatment in both HCT116 and MDA-MB-231 models (Figure 5C, D). In parallel, luciferase-based analysis showed that rAAV-SA significantly increased tumor-cell lysis relative to rAAV-MCS, whereas rAAV-ARMS further increased lysis beyond that achieved with rAAV-SA in both spheroid models (Figure 5E). Consistent with enhanced CAR-T-cell effector function, Granzyme B concentrations in coculture supernatants were increased following rAAV-SA treatment and further elevated with rAAV-ARMS (Figure 5F). Together, these findings demonstrate that incorporation of the IL-15 complex, CXCL10, and IFN-γ enhanced CAR-T-cell effector activity and tumor-cell killing beyond synthetic antigen redirection alone in three-dimensional tumor models.

**Figure 5.**
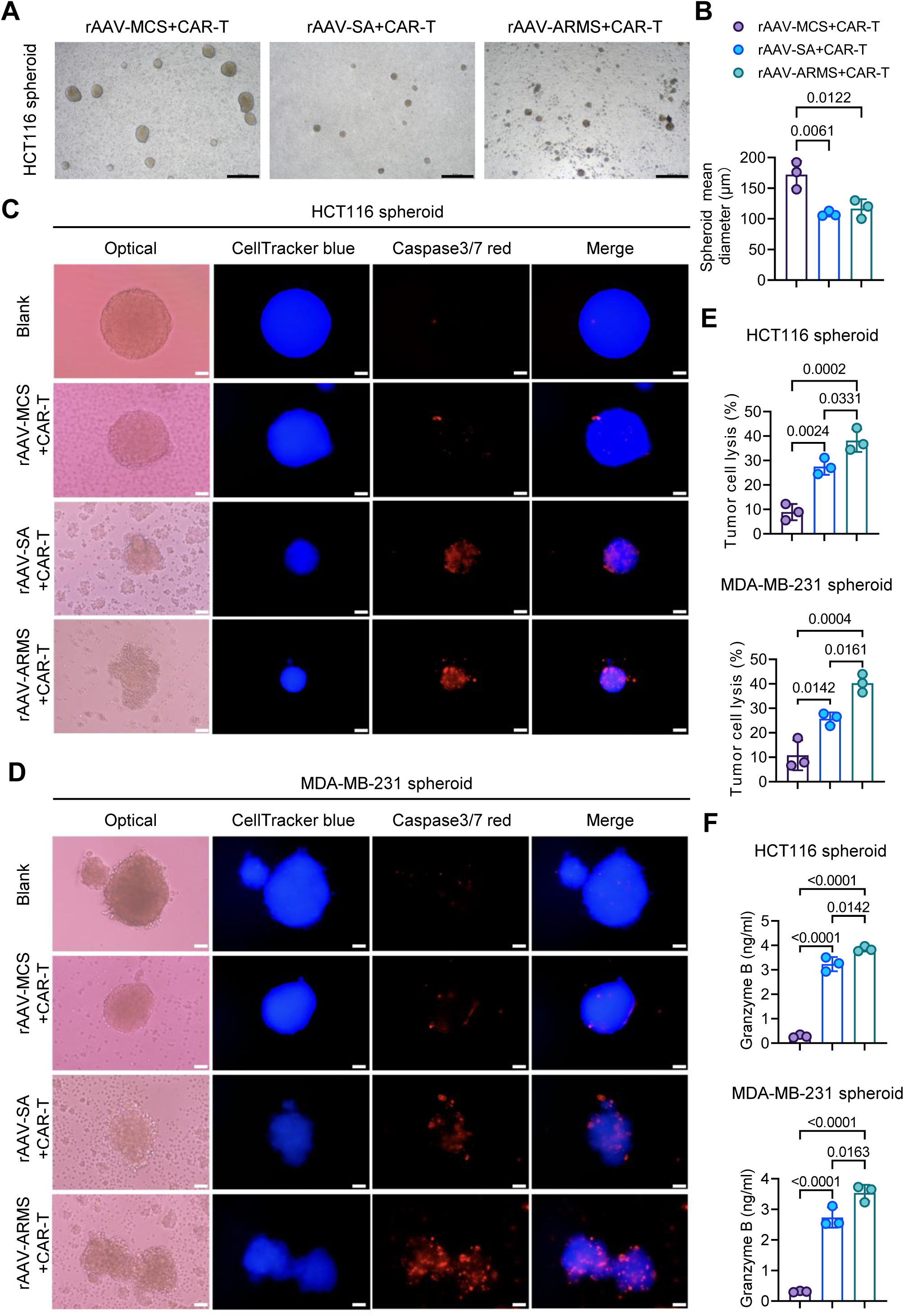
rAAV-ARMS enhances CAR-T-cell cytotoxicity in tumor spheroids. HCT116 and MDA-MB-231 spheroids were treated with rAAV-MCS, rAAV-SA, or rAAV-ARMS for 72 h, followed by CAR-T-cell coculture at an effector-to-target (E:T) ratio of 10:1 for 48 h before endpoint analyses. Blank denotes spheroids treated with neither rAAV nor CAR-T cells. (A) Representative bright-field images of HCT116 spheroids following treatment with rAAV-MCS, rAAV-SA, or rAAV-ARMS and coculture with CAR-T cells. Scale bars, 500 μm. (B) Quantification of mean HCT116 spheroid diameter. (C, D) Representative fluorescence images of HCT116 (C) and MDA-MB-231 (D) spheroids prelabeled with CellTracker Blue before CAR-T-cell coculture and stained with a Caspase-3/7 Red fluorescent reporter at the experimental endpoint. Scale bars, 50 μm. (E) Luciferase-based quantification of tumor-cell lysis (%) in HCT116-Luc and MDA-MB-231-Luc spheroids. (F) Granzyme B concentrations in coculture supernatants measured by ELISA. Data are presented as mean ±SD from three independent experiments. Statistical significance was assessed by one-way ANOVA with Tukey’s test. P values are shown in the figure.

### Multifunctional rAAV-ARMS potentiates systemic CAR-T-cell therapy in a human TNBC model

We next evaluated whether rAAV-ARMS could enhance CAR-T-cell therapy after systemic T-cell administration, a more stringent setting in which circulating CAR-T cells must traffic to and accumulate within tumors. In this design, CXCL10 was incorporated to recruit CXCR3-expressing T cells, while the IL-15 complex and IFN-γ were intended to support T-cell expansion and adhesion within the tumor microenvironment (Figure 6A). We first examined whether rAAV-ARMS retained its multifunctional expression profile in vivo. When tumors reached approximately 100 mm³on day 22, MDA-MB-231 tumor-bearing NCG mice received a single intravenous dose of rAAV-MCS, rAAV-SA, or rAAV-ARMS (1 × 10¹¹vg/mouse), and tumors were analyzed 7 days later (Figure 6B). Both rAAV-SA and rAAV-ARMS induced surface PEPvIII expression in tumors, whereas rAAV-MCS remained at background levels (Figure 6C). In parallel, the IL-15 complex, CXCL10, and IFN-γ were readily detected in tumor homogenates from rAAV-ARMS-treated mice (Figure 6D). Myc immunohistochemistry further confirmed transgene expression in tumors, while no appreciable staining was observed in the heart, liver, spleen, lung, or kidney (Supplementary Figure 10A, B). Notably, the three secreted payloads were undetectable in serum (Supplementary Figure 10C), indicating limited systemic exposure to the secreted payloads despite intravenous rAAV administration.

**Figure 6.**
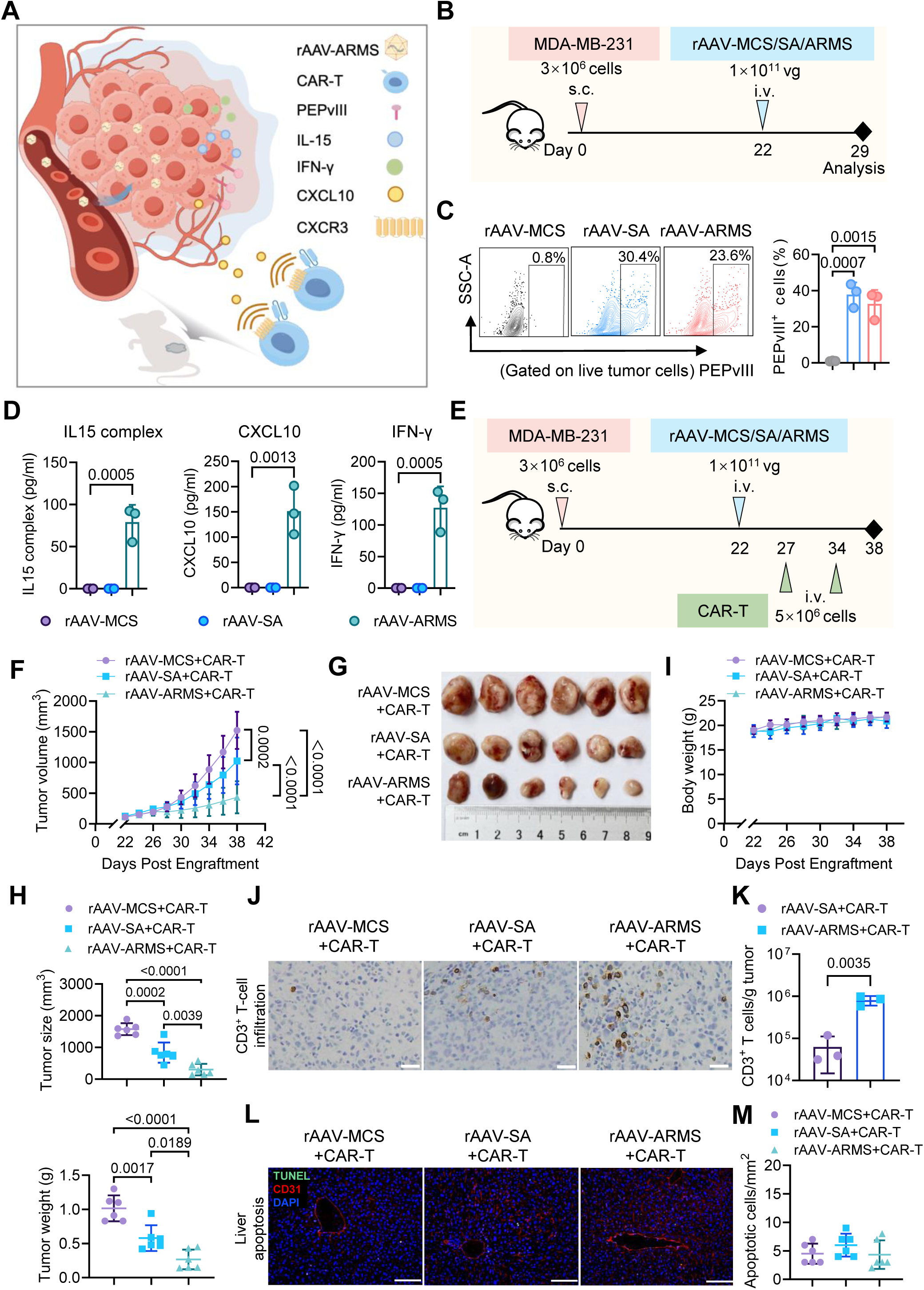
Multifunctional rAAV-ARMS enhances systemic CAR-T-cell therapy in a human TNBC xenograft model. (A) Schematic of the in vivo treatment strategy combining rAAV-ARMS with systemically administered CAR-T cells. (B–D) In vivo characterization of rAAV-ARMS expression in the MDA-MB-231 xenograft model. When tumors reached approximately 100 mm³, mice received a single intravenous dose of rAAV-MCS, rAAV-SA, or rAAV-ARMS (1 ×10¹¹vg/mouse) on day 22, and tumors were collected 7 days later. Experimental design (B), representative flow cytometry plots and quantification of surface PEPvIII expression in dissociated tumor cells (C), and concentrations of the IL-15/IL-15Rα complex, CXCL10, and IFN-γ in tumor homogenates measured by ELISA (D) are shown (n = 3 mice per group). (E–I) Therapeutic evaluation in the MDA-MB-231 xenograft model. When tumors reached approximately 100 mm³, mice received a single intravenous dose of rAAV-MCS, rAAV-SA, or rAAV-ARMS (1 × 10¹¹vg/mouse) on day 22, followed by intravenous CAR-T-cell administration (5 × 10⁶ cells/mouse) on days 27 and 34. Treatment scheme (E), tumor growth curves (F), excised tumors (G), ex vivo tumor volumes and weights measured at the experimental endpoint (H), and body-weight changes (I) are shown (n = 6 mice per group). (J, K) Intratumoral T-cell accumulation in MDA-MB-231 tumors following treatment as shown in (E). Representative CD3 immunohistochemistry images (J) and counts of infiltrating human CD45⁺CD3⁺ T cells per gram of tumor (K) are shown. For flow cytometry, tumors from the rAAV-SA + CAR-T and rAAV-ARMS + CAR-T groups were collected at the experimental endpoint and dissociated into single-cell suspensions (n = 3 mice per group). Scale bar, 25 μm in (J). (L, M) Hepatic apoptosis assessment at the experimental endpoint following treatment as shown in (E). Representative liver sections stained for TUNEL, CD31, and DAPI (L) and quantification of TUNEL-positive cells per mm² (M) are shown. Scale bar, 100 μm in (L). Data are presented as mean ±SD. Statistical significance was assessed using an unpaired two-tailed Student’s t-test for (K), one-way ANOVA followed by Tukey’s multiple-comparisons test for (C), (D), (H), and (M), and two-way ANOVA for (F). P values are shown in the figure.

We then tested the complete strategy using intravenous CAR-T-cell delivery. When tumors reached approximately 100 mm³, MDA-MB-231 tumor-bearing mice received a single intravenous dose of rAAV-MCS, rAAV-SA, or rAAV-ARMS (1 ×10¹¹vg/mouse) on day 22, followed by two intravenous infusions of CAR-T cells (5 × 10⁶ cells/mouse per infusion) on days 27 and 34 (Figure 6E). rAAV-SA combined with CAR-T cells significantly delayed tumor growth compared with rAAV-MCS combined with CAR-T cells, whereas rAAV-ARMS combined with CAR-T cells further suppressed tumor growth (Figure 6F). At the experimental endpoint, tumors from the rAAV-ARMS group were visibly smaller and exhibited significantly reduced ex vivo tumor volume and tumor weight compared with both control groups (Figure 6G, H). Body weight remained stable throughout treatment (Figure 6I). To determine whether the improved therapeutic effect was associated with increased intratumoral T-cell accumulation, we examined T-cell infiltration in tumor tissues. CD3 immunohistochemistry showed greater T-cell accumulation in rAAV-ARMS-treated tumors than in the rAAV-MCS and rAAV-SA groups (Figure 6J). Consistent with this observation, the number of human CD45⁺CD3⁺ T cells per gram of tumor was significantly higher in the rAAV-ARMS group than in the rAAV-SA group (Figure 6K), indicating that the ARMS payload combination was associated with increased intratumoral T-cell accumulation in vivo. Because AAV2 exhibits hepatic tropism, we specifically examined the liver for potential treatment-related injury. TUNEL staining revealed no increase in hepatic apoptosis, including in regions surrounding CD31⁺ vascular structures (Figure 6L, M). All measured serum biochemical values were within their respective reference intervals (Supplementary Figure 11A), and histological examination showed no evident abnormalities in the heart, liver, spleen, lung, or kidney (Supplementary Figure 11B). Together, these findings demonstrate that rAAV-ARMS improves the efficacy of systemically delivered CAR-T cells, accompanied by increased intratumoral T-cell accumulation, while maintaining preferential tumor expression and showing no evident systemic or hepatic toxicity.

## Discussion

In this study, we developed an AAV-based platform that integrates tumor-selective synthetic-antigen redirection with localized delivery of immunomodulatory factors to enhance CAR-T-cell therapy. The platform addresses two major barriers in solid tumors: the limited availability of suitable surface targets and an immunosuppressive tumor microenvironment.^31,32^ We therefore focused on engineering tumor cells as an alternative to further modifying CAR-T cells. We first employed the NF-κB-responsive decoy minimal promoter (DMP) to drive tumor-selective surface display of a synthetic CAR target following rAAV-SA delivery, enabling CAR-T-cell activation and cytotoxicity across multiple tumor cell lines and antitumor activity in human TNBC and colorectal cancer xenograft models. We then incorporated an IL-15/IL-15Rα sushi-domain fusion, CXCL10, and IFN-γ to generate rAAV-ARMS, a multifunctional vector designed to provide local immune support. Relative to antigen delivery alone, rAAV-ARMS enhanced tumor-cell killing in three-dimensional spheroids and improved the efficacy of intravenously administered CAR-T cells in a human TNBC xenograft model. Together, these findings establish a new tumor-centered framework for advancing cellular immunotherapy in solid tumors.

For antigen redirection in solid tumors, both the availability of a recognizable target and the risk of off-tumor recognition must be considered. Heterogeneous expression or loss of tumor antigens can compromise recognition and contribute to immune escape and therapeutic resistance.^30,33^ Notably, a recent elegant study demonstrated that MAPK inhibition restores TSHR expression in aggressive thyroid cancers, increasing target antigen density and sensitizing tumors to TSHR-directed CAR-T-cell therapy.^34^ This study highlights the therapeutic potential of modifying tumor-intrinsic biology to enhance susceptibility to CAR-T-cell recognition and killing. Similarly, we exploited aberrantly activated NF-κB signaling to introduce a synthetic CAR target, enabling the recognition of tumor cells lacking the corresponding endogenous antigen. However, improving target availability must be balanced against the risk of off-tumor recognition. Even clinically established CD19-directed CAR-T-cell therapy can deplete normal CD19-expressing B cells, resulting in B-cell aplasia and hypogammaglobulinemia that may compromise humoral immunity.^35^ Although logic-gated CAR designs can improve tumor discrimination by integrating multiple antigen inputs,^36–38^ they require more complex receptor engineering and remain dependent on specific combinations of endogenous tumor antigens. In contrast, we selected PEPvIII, a peptide derived from the tumor-specific EGFRvIII junctional epitope, as a model antigen because this epitope is absent from normal tissues and can be recognized by EGFRvIII-directed CAR-T cells.^30,39,40^ Combining this synthetic antigen with DMP-mediated tumor-responsive expression may reduce unintended antigen display in normal tissues and the associated risk of off-tumor recognition. In vitro, flow cytometry detected synthetic-antigen surface expression in the tumor cell lines examined, whereas signals in non-tumor cells remained near background levels. Despite broad rAAV biodistribution in vivo, synthetic-antigen expression was predominantly detected in tumor tissues, consistent with DMP-mediated transcriptional control. Functional assays and TNBC and colorectal cancer xenograft studies further demonstrated that the introduced antigen supported CAR-T-cell activation, degranulation, cytotoxicity, and antitumor activity. Together, these findings highlight the potential of combining rAAV-mediated delivery with tumor-responsive transcriptional control to establish synthetic CAR targets while limiting their expression in normal tissues.

In parallel with synthetic CAR-target installation, we sought to address the immunosuppressive TME by extending tumor-cell engineering to localized immune support. Through NF-κB-responsive expression, rAAV-ARMS programs tumor cells as local sources of immunomodulatory factors, establishing a tumor-side armoring strategy that couples synthetic-antigen display with immune support. Recent clinical evidence indicates that IL-15 armoring can enhance CAR-T-cell expansion, intratumoral persistence, and antitumor activity in solid tumors.^41^ Consistent with the proliferative activity of IL-15/IL-15Rα signaling,^42^ rAAV-ARMS-conditioned medium increased Ki67 expression in CD8⁺ T cells. CXCL10 was incorporated to facilitate T-cell trafficking. In spheroid cocultures, bright-field imaging suggested increased cell accumulation around rAAV-ARMS-treated tumor spheroids. In vivo, rAAV-ARMS substantially increased intratumoral T-cell abundance following systemic CAR-T-cell administration in a TNBC model, consistent with recent evidence that tumor-localized CXCL10 promotes CAR-T-cell infiltration and tumor control.^15^ rAAV-ARMS also increased tumor-cell ICAM-1 expression, a change that may favor CAR-T-cell–tumor cell interactions given the established role of IFN-γ in upregulating ICAM-1 and reinforcing ICAM-1–LFA-1-dependent adhesion.^16^ Although the individual contribution of each effector was not dissected in the present study, their combined delivery produced a functional gain beyond antigen redirection alone, with greater tumor-cell killing and granzyme B release in three-dimensional spheroids and improved tumor control following intravenous CAR-T-cell administration. Given the substantial hepatic exposure associated with systemic AAV delivery,^43^ we further evaluated potential off-target and systemic toxicity. Despite vector distribution beyond the tumor, circulating IL-15 complex, CXCL10, and IFN-γ remained below the assay detection limits, while hepatic TUNEL staining, serum ALT, AST, and ALP levels, and histopathological examination of major organs revealed no overt treatment-related abnormalities in the TNBC model. Together, these findings support tumor-side armoring as a complementary approach to enhancing CAR-T-cell therapy through localized immune support, with limited detectable peripheral cytokine exposure and no overt systemic toxicity under the conditions examined.

While this study establishes a promising framework for tumor-directed immune engineering, several translational considerations require further evaluation. One safety concern is whether NF-κB-responsive expression could be induced in activated immune cells,^44^ potentially resulting in ectopic synthetic-antigen display and subsequent CAR-T-cell–mediated fratricide or bystander cytotoxicity. To examine this possibility, we assessed rAAV-DMP-ZsGreen reporter expression in unstimulated and activated primary T cells. Reporter signals remained at background levels in both T-cell populations, whereas MDA-MB-231 cells showed readily detectable expression. This observation is consistent with the relatively low permissiveness of primary human T cells to unmodified AAV2 reported previously,^45^ which may provide an additional layer of protection against unintended antigen display in immune cells. In addition to these considerations of expression specificity, rAAV-ARMS may exert broader immunomodulatory effects within the TME. IFN-γ, for example, can enhance antigen presentation and potentially promote a more inflamed tumor state.^46,47^ Future studies using CD34⁺ HSC-humanized models will be important to evaluate transgene expression across human immune-cell subsets and explore the potential of rAAV-ARMS to reshape the tumor immune microenvironment.

Humoral immunity to AAV represents another translational challenge. Pre-existing anti-capsid neutralizing antibodies are common and can impair vector transduction, whereas vector-induced antibodies can restrict subsequent dosing.^48^ Encouragingly, recent capsid-engineering approaches have generated serologically distinct AAV variants capable of evading neutralization and enabling redosing.^49^ In parallel, imlifidase-mediated IgG cleavage has restored AAV transduction in seropositive preclinical models and was recently reported to enable AAV gene therapy in a seropositive patient.^50,51^ These advances may support repeated rAAV-ARMS administration to sustain synthetic-antigen availability during treatment, potentially reducing the risk of antigen-loss-mediated escape.

Recent advances in in vivo CAR-T-cell engineering offer an opportunity to further extend the therapeutic potential of rAAV-ARMS. Studies using targeted nanocarriers,^52^ lipid nanoparticles,^53^ engineered lentiviral vectors,^54^ and AAV-assisted genome-editing systems have demonstrated the feasibility of generating functional CAR-T cells directly in vivo.^55^ Building on these advances, rAAV-ARMS could be combined with an engineered T-cell-targeted AAV or Lentivirus carrying the CAR expression cassette used in this study to form a dual-virus cocktail for single-administration treatment. By integrating tumor-preferential synthetic-antigen expression, localized immune support, and in vivo CAR-T-cell generation, this strategy could bypass ex vivo CAR-T-cell manufacturing and pave the way for an off-the-shelf therapeutic modality for solid tumors.

In summary, rAAV-ARMS establishes a tumor-programmed strategy for target-redirected universal CAR-T-cell therapy in solid tumors. Through NF-κB-gated transgene expression, it installs a shared synthetic antigen across distinct tumor types, shifting the basis of universality from adapting CARs to endogenous tumor antigens to making diverse tumors recognizable by a single CAR design. Its modular architecture further integrates antigen redirection with adaptable immunomodulatory payloads and may be extended to CAR-NK cells, CAR-macrophages, and compatible bispecific engagers. This strategy redefines the path toward universal CAR targeting: instead of searching for a common antigen across cancers, it creates one.

## Materials and Methods

### Cell lines and cell culture

The cell lines used in this study included HEK293T, MDA-MB-468, MDA-MB-231, MCF-7, PANC-1, HCT116, MCF-12A, MRC-5, GES-1, HL7702, and NIH-3T3 cells. HEK293T, MDA-MB-468, MDA-MB-231, MCF-7, PANC-1, HCT116, and NIH-3T3 cells were cultured in DMEM (Gibco), whereas MCF-12A, MRC-5, GES-1, and HL7702 cells were maintained in RPMI 1640 medium (Gibco). All culture media were supplemented with 10% fetal bovine serum, 100 U/mL penicillin, and 100 μg/mL streptomycin (Gibco). Cells were maintained at 37°C in a humidified incubator containing 5% CO₂.

### DNA constructs

The NF-κB-responsive decoy minimal promoter (DMP), previously developed by our group, consists of five tandem κB-binding sites followed by a minimal promoter.^56^ The previously constructed pAAV-DMP-hCD20 plasmid was used as the backbone for generation of the DMP-driven expression vectors.^28^ The synthetic antigen (SA) cassette, comprising an Igκ leader sequence, PEPvIII (a peptide derived from the EGFRvIII junctional epitope), a Myc tag, a (G4S)4 flexible linker, and the transmembrane domain of platelet-derived growth factor receptor (PDGFR), was chemically synthesized by GenScript. The SA fragment was amplified using primers containing EcoRV and BamHI restriction sites and inserted into the pAAV-DMP backbone to replace the hCD20 coding sequence, generating pAAV-DMP-SA. The ZsGreen coding sequence was also chemically synthesized by GenScript, amplified using EcoRV- and BamHI-containing primers, and inserted into the same backbone to generate pAAV-DMP-ZsGreen. Based on the pAAV-DMP-SA backbone, DNA fragments encoding an IL-15Rα sushi-domain/IL-15 fusion, CXCL10, and IFN-γ were chemically synthesized by GenScript and incorporated to generate pAAV-ARMS. Each secreted effector was driven by an independent DMP and terminated by its own polyadenylation signal, resulting in a single vector integrating the synthetic antigen module with three secreted effector cassettes.

An anti-EGFRvIII CAR was generated using pSLCAR-CD19-BBz (Addgene) as the backbone. A DNA fragment encoding the CD8α signal peptide together with a previously reported EGFRvIII-specific scFv was chemically synthesized by GenScript,^30^ amplified using primers containing BamHI and KpnI restriction sites, and inserted into the CAR backbone to replace the corresponding CD8α signal peptide–FMC63 scFv region. The remaining CAR components, including the CD8 hinge, CD28 transmembrane domain, 4-1BB/CD3ζ intracellular signaling domains, and P2A-linked EGFP reporter, were retained. Primer sequences are listed in Supplementary Table 1.

### Recombinant AAV production and titration

HEK293T cells were seeded at 5 × 10⁶ cells per flask and cultured overnight before cotransfection with pHelper, pAAV-RC, and the indicated pAAV transfer plasmid using Lipofectamine 3000 (Thermo Fisher Scientific, Cat. No. L3000015). The pAAV-RC plasmid provided the AAV2 rep and cap genes required for rAAV2 packaging. At 72 h post-transfection, cells and culture supernatants were harvested and subjected to three freeze–thaw cycles for viral particle release. The crude lysate was extracted with chloroform, adjusted to 1 M NaCl, and clarified by centrifugation. rAAV particles in the clarified supernatant were precipitated with polyethylene glycol 8000 (Sigma-Aldrich, Cat. No. 89510), resuspended in PBS, and treated with DNase and RNase (1 μg/mL; Sangon Biotech) to remove residual nucleic acids, followed by a second chloroform extraction. Vector genome titers were determined by quantitative PCR using primers listed in Supplementary Table 2 and expressed as vector genomes per milliliter (vg/mL). The resulting vector preparations were designated rAAV-MCS, rAAV-SA, rAAV-ARMS, and rAAV-DMP-ZsGreen.

### Lentivirus production and concentration

HEK293T cells were cotransfected with the anti-EGFRvIII CAR transfer plasmid, psPAX2, and pMD2.G using Lipofectamine 3000. At 48 h after transfection, culture supernatants containing lentiviral particles were collected and concentrated using the Universal Virus Concentration Kit (Beyotime, Cat. No. C2901S) according to the manufacturer’s instructions.

### Quantitative PCR

Total RNA was extracted from cultured cells and tissues using TRIzol reagent (Invitrogen, Cat. No. 15596026), and cDNA was synthesized using the PrimeScript RT Reagent Kit with gDNA Eraser (Takara, Cat. No. RR047A). Genomic DNA was isolated from tissues using the TIANamp Genomic DNA Kit (TIANGEN, Cat. No. DP304-02). Quantitative PCR was performed on an ABI StepOne Plus Real-Time PCR System (Applied Biosystems) using FastStart Universal SYBR Green Master (ROX) (Roche, Cat. No. 04913850001). Relative mRNA expression was calculated using the 2^^−ΔCt^ method with GAPDH as the internal reference. Primer sequences are listed in Supplementary Table 2.

### Assessment of synthetic antigen and ZsGreen expression

For assessment of synthetic antigen expression, cells were transfected with pAAV-MCS or pAAV-DMP-SA using Lipofectamine 3000. After 72 h, cells were stained with PE-conjugated anti-EGFRvIII antibody (clone DH8.3.Rec, BioLegend), and cell-surface PEPvIII expression was quantified by flow cytometry. Freshly isolated T cells, T cells activated with CD3/CD28 for 48 h, and MDA-MB-231 cells were treated with rAAV-MCS or rAAV-DMP-ZsGreen at an MOI of 1 × 10⁵ vg/cell, followed by flow-cytometric analysis of ZsGreen fluorescence 72 h later.

### Generation of CAR-T cells

Peripheral blood samples were collected with written informed consent under a protocol approved by the Committee for Ethical Review of Research Involving Human Subjects of Nanjing Hospital of Chinese Medicine Affiliated to Nanjing University of Chinese Medicine (approval no. KY2025052). PBMCs were isolated by density-gradient centrifugation using Lymphoprep (STEMCELL Technologies, Cat. No. 18061). CD3⁺ T cells were purified using the EasySep Human T Cell Isolation Kit (STEMCELL Technologies, Cat. No. 17951) according to the manufacturer’s instructions. Purified T cells were resuspended at 1 × 10⁶ cells/mL in ImmunoCult-XF T Cell Expansion Medium supplemented with 10 ng/mL recombinant human IL-2 (STEMCELL Technologies, Cat. No. 78036) and activated with ImmunoCult Human CD3/CD28 T Cell Activator at 25 μL/mL. At 24 h after activation, T cells were transduced with lentivirus encoding the anti-EGFRvIII CAR at an MOI of 5. After 24 h of transduction, the viral supernatant was replaced with fresh ImmunoCult-XF medium containing 10 ng/mL IL-2, and cells were expanded before subsequent experiments. Transduction efficiency was assessed by EGFP expression. CAR surface expression was independently evaluated using a fluorescent PEPvIII probe generated by pre-complexing N-terminally biotinylated PEPvIII (LEEKKGNYVVTDHGS; GenScript) with PE-conjugated streptavidin (BioLegend, Cat. No. 405203), followed by flow-cytometric analysis.

### CAR-T-cell functional assays

Before functional assays, CAR-T cells were washed twice with fresh culture medium to remove residual exogenous IL-2 and CD3/CD28 activation reagents. MDA-MB-231 and HCT116 cells were transfected with pAAV-MCS or pAAV-DMP-SA for 72 h and subsequently cocultured with CAR-T cells at an effector-to-target (E:T) ratio of 5:1 for 18 h. IL-2, IFN-γ, and TNF-α in culture supernatants were quantified using Human IL-2 ELISA Kit (Elabscience, Cat. No. E-EL-H0099), Human IFN-γ ELISA Kit (Elabscience, Cat. No. E-EL-H0108), and Human TNF-α ELISA Kit (Elabscience, Cat. No. E-EL-H0109), respectively.

For assessment of CAR-dependent activation, CAR-T cells cocultured with MDA-MB-231 target cells were analyzed for CD69 and CD25 expression within CAR⁺ and CAR⁻ T-cell populations identified by EGFP expression, using anti-human CD69-APC (clone FN50, BioLegend) and anti-human CD25-APC (clone BC96, BioLegend), respectively. CAR-T-cell degranulation following coculture with MDA-MB-231 target cells was evaluated by CD107a surface expression within CAR⁺CD8⁺ T cells using anti-human CD107a-APC (clone H4A3, BioLegend) and anti-human CD8-PE (clone HIT8a, BioLegend). The anti-CD107a antibody was added 4 h before the end of the 18-h coculture.

For direct assessment of tumor-cell killing, MDA-MB-231, MCF-7, PANC-1, and HCT116 cells were transfected with pAAV-MCS or pAAV-DMP-SA for 72 h and labeled with CellTrace Far Red (Thermo Fisher Scientific, Cat. No. C34564). Labeled target cells were then cocultured with CAR-T cells at an E:T ratio of 5:1 for 18 h. Following coculture, cells were stained with propidium iodide (PI). Target cells were identified as CellTrace Far Red⁺ events, and cytotoxicity was quantified as the percentage of PI⁺ cells within the labeled tumor-cell population.

### Tumor spheroid culture and CAR-T-cell cytotoxicity

MDA-MB-231 and HCT116 cells were seeded at 1 ×10³cells per well in ultra-low-attachment 96-well plates (Corning) and cultured under serum-free suspension conditions. HCT116 spheroids were cultured for 48 h and MDA-MB-231 spheroids for 7 days before treatment with rAAV-MCS, rAAV-SA, or rAAV-ARMS at 1 × 10⁸ vg/well. At 72 h after rAAV treatment, spheroids were dissociated with TrypLE Express (Thermo Fisher Scientific, Cat. No. 12604013) for flow-cytometric analysis of surface PEPvIII and ICAM-1 expression. Concentrations of the IL-15/IL-15Rα complex in culture supernatants were measured using a Human IL-15/IL-15Rα Complex DuoSet ELISA (R&D Systems, Cat. No. DY6924), whereas CXCL10 and IFN-γ were quantified using a Human IP-10/CXCL10 ELISA Kit (Elabscience, Cat. No. E-EL-H0050) and Human IFN-γ ELISA Kit (Elabscience, Cat. No. E-EL-H0108), respectively. To assess the biological activity of secreted IL-15, primary human T cells were cultured for 48 h with conditioned media collected from the indicated spheroids. Cells were subsequently stained with anti-human CD8-PE (clone HIT8a, BioLegend), fixed and permeabilized using the FOXP3 Fix/Perm Buffer Set (BioLegend, Cat. No. 421403), and stained with anti-Ki67-APC (clone Ki-67, BioLegend) for flow-cytometric analysis.

To assess whether rAAV treatment alone affected spheroid growth or viability, MDA-MB-231 and HCT116 spheroids were treated with PBS, rAAV-MCS, rAAV-SA, or rAAV-ARMS in the absence of T cells. After 72 h, spheroid morphology and diameter were evaluated by bright-field microscopy, and viability was assessed using the Beyo3D Calcein/PI/Hoechst Cell Viability Assay Kit (Beyotime, Cat. No. C1375S). In parallel, viability of luciferase-expressing HCT116 and MDA-MB-231 spheroids was quantified using the Pierce Firefly Luciferase Glow Assay Kit (Thermo Fisher Scientific, Cat. No. 16176).

For CAR-T-cell cytotoxicity assays, spheroids were treated with rAAV-MCS, rAAV-SA, or rAAV-ARMS for 72 h before coculture with CAR-T cells at an E:T ratio of 10:1 for an additional 48 h. Spheroids treated with neither rAAV nor CAR-T cells served as blank controls. Spheroid diameter was quantified from bright-field images using ImageJ. For fluorescence-based assessment of tumor-cell apoptosis, spheroids were prelabeled with CellTracker Blue according to the manufacturer’s instructions before coculture, and caspase-3/7 activity was detected using CellEvent Caspase-3/7 Red Detection Reagent (Thermo Fisher Scientific, Cat. No. C10730). Tumor-cell killing in HCT116-Luc and MDA-MB-231-Luc spheroids was further quantified by firefly luciferase activity. Tumor-cell lysis was calculated relative to the blank control as [1 − (luminescence of the experimental group/luminescence of the blank control)] × 100%. Granzyme B was quantified using a Human Granzyme B ELISA Kit (Elabscience, Cat. No. E-EL-H1617).

### Animal models

All animal experiments were conducted in accordance with protocols approved by the Animal Experimentation and Ethics Committee of Southeast University. Six-week-old female NOD CRISPR Prkdc Il2rγ (NCG) triple-immunodeficient mice were obtained from GemPharmatech Co., Ltd. MDA-MB-231 (3 × 10⁶ cells/mouse) or HCT116 (5 × 10⁶ cells/mouse) cells were implanted subcutaneously to establish xenograft models. Tumor volume was calculated as V = ab²/2, where a and b represent the longest and shortest tumor diameters, respectively. Tumor growth and body weight were monitored throughout therapeutic studies. Mice were euthanized when tumor volume reached 2,000 mm³or at the predefined experimental endpoint. At therapeutic endpoints, serum was collected for biochemical analysis, and the heart, liver, spleen, lung, and kidney were collected for hematoxylin and eosin (H&E) staining. For flow-cytometric analyses, freshly excised tumors were mechanically dissociated using a gentleMACS Dissociator (Miltenyi Biotec), followed by filtration through 70-μm cell strainers to obtain single-cell suspensions.

For in vivo evaluation of synthetic antigen expression, MDA-MB-231 tumor-bearing mice received a single intravenous dose of rAAV-MCS or rAAV-SA (5 × 10¹⁰ vg/mouse) on day 25, when tumors reached approximately 100 mm³. Tumors and major organs were collected 7 days after rAAV administration (n = 3 mice per group). A portion of each tumor was processed into a single-cell suspension as described above for flow-cytometric analysis of surface PEPvIII expression. Genomic DNA and total RNA were extracted from the heart, liver, spleen, lung, kidney, and tumor tissues for quantification of rAAV DNA abundance and PEPvIII mRNA expression, respectively. Myc-tag immunohistochemistry was performed on paraffin-embedded tumor and major-organ sections using an anti-MYC tag antibody (Proteintech, Cat. No. 16286-1-AP) to assess transgene expression and tissue distribution.

For therapeutic evaluation in the MDA-MB-231 model, when tumors reached approximately 100 mm³, tumor-bearing mice received PBS, rAAV-MCS, or rAAV-SA (5 × 10¹⁰ vg/mouse per rAAV injection) intravenously on days 25, 27, and 29 after tumor implantation, followed by intratumoral administration of CAR-T cells (5 × 10⁶ cells/mouse) on day 31 (n = 7 mice per group). At the experimental endpoint, tumors were excised and weighed.

To assess therapeutic efficacy in the HCT116 model, when tumors reached approximately 100 mm³, tumor-bearing mice received PBS, rAAV-MCS, or rAAV-SA (5 × 10¹⁰ vg/mouse per rAAV injection) intravenously on days 5 and 7, followed by intratumoral CAR-T-cell administration (5 × 10⁶ cells/mouse per injection) on days 9 and 13 (n = 6 mice per group). At the experimental endpoint, tumors and major organs were collected for downstream analyses. Genomic DNA and total RNA were extracted from the heart, liver, spleen, lung, kidney, and tumor tissues for quantification of rAAV DNA abundance and PEPvIII mRNA expression, respectively, and Myc-tag immunohistochemistry was performed on tumor and major-organ sections as described above.

For short-term analysis of intratumoral T-cell activation, when tumors reached approximately 100 mm³, HCT116 tumor-bearing mice received rAAV-MCS or rAAV-SA (5 × 10¹⁰ vg/mouse per injection) intravenously on days 5 and 7, followed by a single intratumoral CAR-T-cell injection (5 × 10⁶ cells/mouse) on day 9. Tumors were collected 2 days later, processed into single-cell suspensions, and analyzed by flow cytometry for CD69 expression in CD3⁺ T cells and CD25 expression in CAR⁺ T cells (n = 3 mice per group).

For evaluation of rAAV-ARMS expression in the MDA-MB-231 model, when tumors reached approximately 100 mm³on day 22, mice received a single intravenous dose of rAAV-MCS, rAAV-SA, or rAAV-ARMS (1 × 10¹¹vg/mouse). Tumors and major organs were collected 7 days after rAAV administration (n = 3 mice per group). Dissociated tumor cells were analyzed by flow cytometry for surface PEPvIII expression, and concentrations of the IL-15/IL-15Rα complex, CXCL10, and IFN-γ in tumor homogenates and serum were measured by ELISA. Myc-tag immunohistochemistry was performed on tumor and major-organ sections as described above.

For therapeutic studies, when tumors reached approximately 100 mm³, mice received a single intravenous dose of rAAV-MCS, rAAV-SA, or rAAV-ARMS (1 × 10¹¹vg/mouse) on day 22 after tumor implantation, followed by intravenous administration of CAR-T cells (5 × 10⁶ cells/mouse per infusion) on days 27 and 34 (n = 6 mice per group). The study was terminated on day 38 when tumor volumes in some mice reached or exceeded the humane endpoint of approximately 2,000 mm³. Tumors, major organs, and serum were collected for downstream analyses. Excised tumors were measured for ex vivo tumor volume and subsequently weighed. Intratumoral T-cell accumulation was assessed by CD3 immunohistochemistry using an anti-human CD3 antibody (Proteintech, Cat. No. 60181-1) and by flow cytometry. For flow-cytometric quantification, tumors from three mice per group in the rAAV-SA combined with CAR-T cells and rAAV-ARMS combined with CAR-T cells groups were analyzed. Total viable cells in dissociated tumors were counted using an automated cell counter. Human CD45⁺CD3⁺ T-cell counts were calculated from their frequency among viable singlets and normalized to tumor weight (cells/g).

Hepatic apoptosis was evaluated using a CF488 TUNEL Cell Apoptosis Detection Kit (Servicebio, Cat. No. G1504-50T), together with CD31 immunofluorescence using an anti-CD31 mouse recombinant monoclonal antibody (Servicebio, Cat. No. GB15063-100) and DAPI nuclear counterstaining (Servicebio, Cat. No. G1012). TUNEL-positive cells were quantified per mm²of liver tissue. Serum ALT, AST, ALP, BUN, creatinine, and uric acid levels were measured for biochemical safety assessment.

### Flow cytometry

Flow-cytometric data were acquired using a NovoCyte 2000R flow cytometer (Agilent Technologies) and analyzed using FlowJo v10.8 software. Antibodies used for flow-cytometric analyses are listed in Supplementary Table 3.

### Statistical analysis

Statistical analyses were performed using GraphPad Prism 8.0. Data are presented as mean ± SD. Comparisons between two groups were performed using unpaired two-tailed Student’s t-tests. Comparisons among three or more groups were performed using one-way ANOVA with Tukey’s multiple-comparisons test or two-way ANOVA with appropriate multiple-comparisons testing, as indicated in the figure legends. A P value < 0.05 was considered statistically significant.

## Data availability

The main data supporting the findings of this study are available within the Article and the Supplementary Information or from the corresponding author upon reasonable request. The source data underlying Figs. 1B and 1C; 2C–2F and 2H; 3B, 3E, 3G, 3H, 3J, 3L, and 3M; 4C, 4D, 4F, and 4H; 5B, 5E, and 5F; and 6C, 6D, 6F, 6H, 6I, 6K, and 6M, as well as Supplementary Figs. 3B; 4B–4E; 5A and 5B; 6A–6C; 7A and 7B; 8B; 9B, 9E, and 9F; and 11A, are provided with the paper as a Source Data file.

## Acknowledgments

The graphical abstract and schematic illustrations in Figures 1A, 2A, 4A, and 6A were prepared using Figdraw (www.figdraw.com).

## Author contributions

Y.K. under direction of J.W. designed and performed experiments, analyzed the data, and wrote first version of manuscript. Y.W., Y.Z., A.Y, X.W., J.X., and L.W performed part of experiments. J.W., F.Z., and Y.S. designed experiments and supervised the study. Y.Z. and J.W. wrote the manuscript. All authors contributed to the critical revision of the manuscript and approved the final version.

## Ethics approval and consent to participate

Human peripheral blood collection was approved by the Committee for Ethical Review of Research Involving Human Subjects of Nanjing Hospital of Chinese Medicine Affiliated to Nanjing University of Chinese Medicine (approval no. KY2025052), and written informed consent was obtained from all participants. All animal experiments were approved by the Animal Experimentation and Ethics Committee of Southeast University.

