## Supplemental Table 1-3 and Supplemental Figures 1-11 for "Tumor-responsive AAV arms cancer cells to enhance CAR-T cell therapy in solid tumors"

**Supplementary Table 1. Primer sequences used for plasmid construction**

| Name | Primer Sequence (5’-3’) |
|---|---|
| SA-F | GATATCGCCACCATGGAGACAGACAC |
| SA-R | GGATCCCTAACGTGGCTTCTTCT |
| ZsGreen-F | GATATCGCCACCATGGCCCAGTCC |
| ZsGreen-R | GGATCCTTAGGGCAAGGCGGA |
| EGFRvIII-scFv-F | GGATCCGCCACCATGGCCCTGCCGG |
| EGFRvIII-scFv-R | GGTACCCTTGATCTCCACCTTG |

**Supplementary Table 2. Primer sequences used for qPCR**

| Name | Primer Sequence (5’-3’) |
|---|---|
| Human RELA-F | CCTGGAGCAAGGACTCAGCA |
| Human RELA-R | ATGCACAGCAGGACAATGGG |
| Human GAPDH-F | ATTTTGGAGGGATCTCGCTCC |
| Human GAPDH-R | CTCCCTCTTCCGTTCCAGTTT |
| Mouse RELA-F | AGGCTTCTGGGCCTTATGTG |
| Mouse RELA-R | TGCTTCTCTCGCCAGGAATAC |
| Mouse GAPDH-F | TCACCACCATGGAGAAGGC |
| Mouse GAPDH-R | GCTAAGCAGTTGGTGGTGCA |
| PEPvIII-F | GTGACCTGGAAGAGAAGAAG |
| PEPvIII-R | TTCGTCGACCTGCAGATCG |
| AAV-F | TGCATGACCAGGCTCAGCTA |
| AAV-R | GACAGGGAAGGGAGCAGTG |
| Mouse gGAPDH-F | GGCCACGCTAATCTCATTTT |
| Mouse gGAPDH-R | AAGGCGGAGTTACCAGAGGT |
| Human gGAPDH-F | CTGACTCAGCCCTGCAAAG |
| Human gGAPDH-R | CCTGCCTTCCTCACCTGAT |

**Supplementary Table 3. Flow cytometry staining panel**

| Marker | Tag | Manufacturer | Clone |
|---|---|---|---|
| Viability staining | | | |
| 7-AAD | — | BioLegend | — |
| Fc receptor blocking | | | |
| Human TruStain FcX | — | BioLegend | — |
| Mouse CD16/32 | — | BioLegend | S17011E |
| Surface staining | | | |
| Anti-EGFRvIII | PE | BioLegend | DH8.3.Rec |
| Human CD3 | PerCP | BioLegend | SK7 |
| Human CD3 | PE | BioLegend | SK7 |
| Human CD4 | APC | BioLegend | RPA-T4 |
| Human CD8 | PE | BioLegend | HIT8a |
| Human CD25 | APC | BioLegend | BC96 |
| Human CD69 | APC | BioLegend | FN50 |
| Human CD107a | APC | BioLegend | H4A3 |
| Human ICAM-1 | FITC | BioLegend | HCD54 |
| Human CD45 | APC | BioLegend | 2D1 |
| Intracellular/nuclear staining[†] | | | |
| Ki-67 | APC | BioLegend | Ki-67 |

† Cells were fixed and permeabilized using the FOXP3 Fix/Perm Buffer Set (BioLegend).

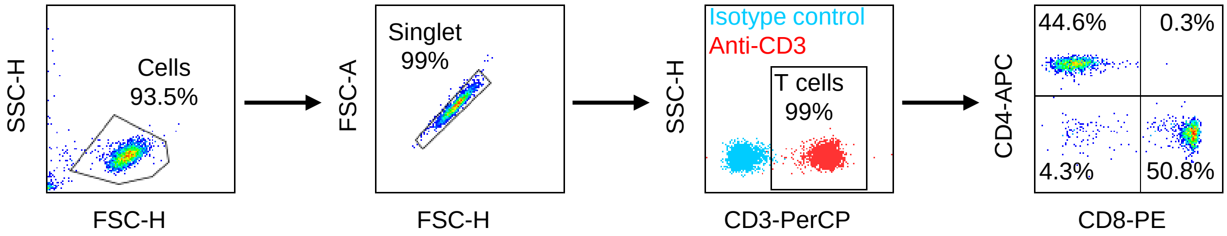


**Supplementary Figure 1. Purity assessment of peripheral blood-derived T cells prior to CAR-T cell generation.** Representative flow cytometry plots showing the gating strategy for evaluating T-cell purity. After exclusion of debris and doublets, T cells were identified as $CD3^+$ cells using anti-CD3 staining, with an isotype control used to define background fluorescence. The isolated T-cell population exhibited approximately 99% $CD3^+$ purity. CD4/CD8 profiling of the $CD3^+$ population showed 44.6% $CD4^+CD8^-$ and 50.8% $CD4^-CD8^+$ T cells. The purified T cells were subsequently used for CAR-T cell generation.

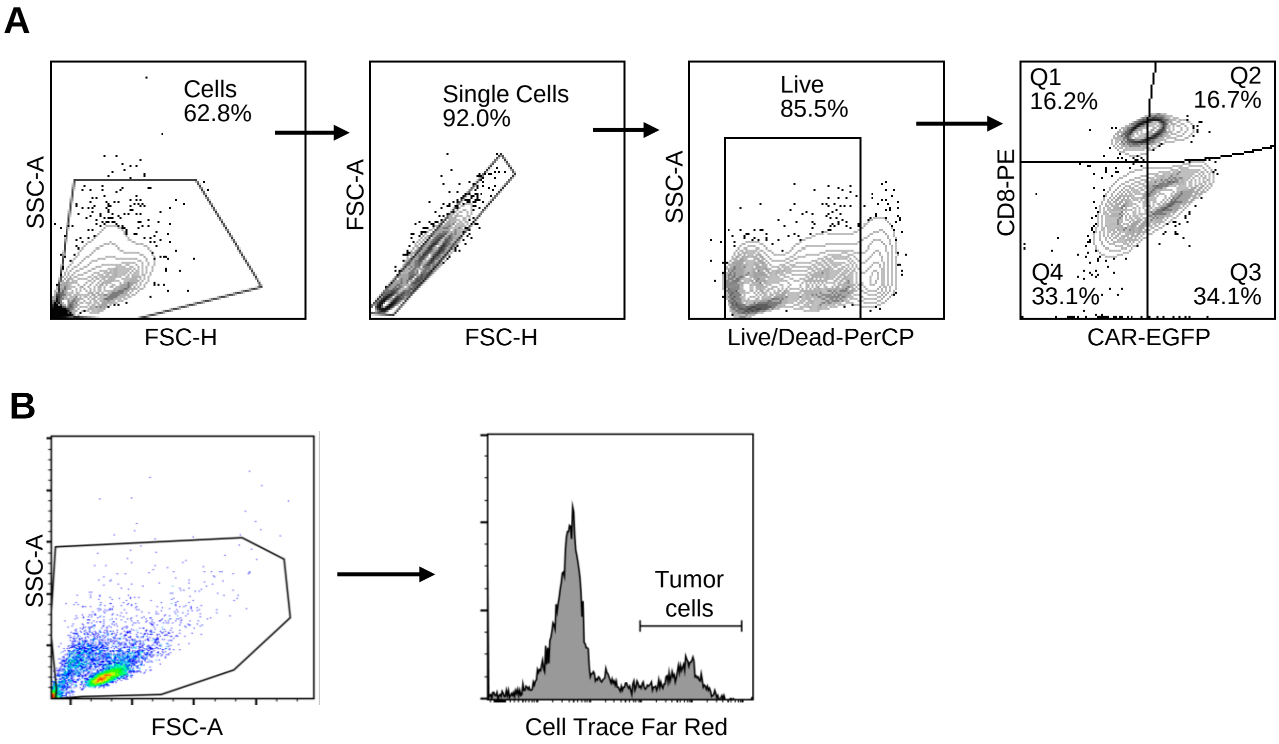


**Supplementary Figure 2. Flow cytometry gating strategies for CAR-T-cell functional assays.** (A) Representative gating strategy for identification of CAR$^+$CD8$^+$ T cells. Cells were sequentially gated for cells, singlets, and live cells, followed by identification of CAR$^+$CD8$^+$ T cells based on EGFP and CD8 expression. (B) Representative gating strategy for identification of target tumor cells in the cytotoxicity assay. CellTrace Far Red-labeled tumor cells were identified as the CellTrace Far Red$^+$ population for subsequent PI-based assessment of tumor-cell death.

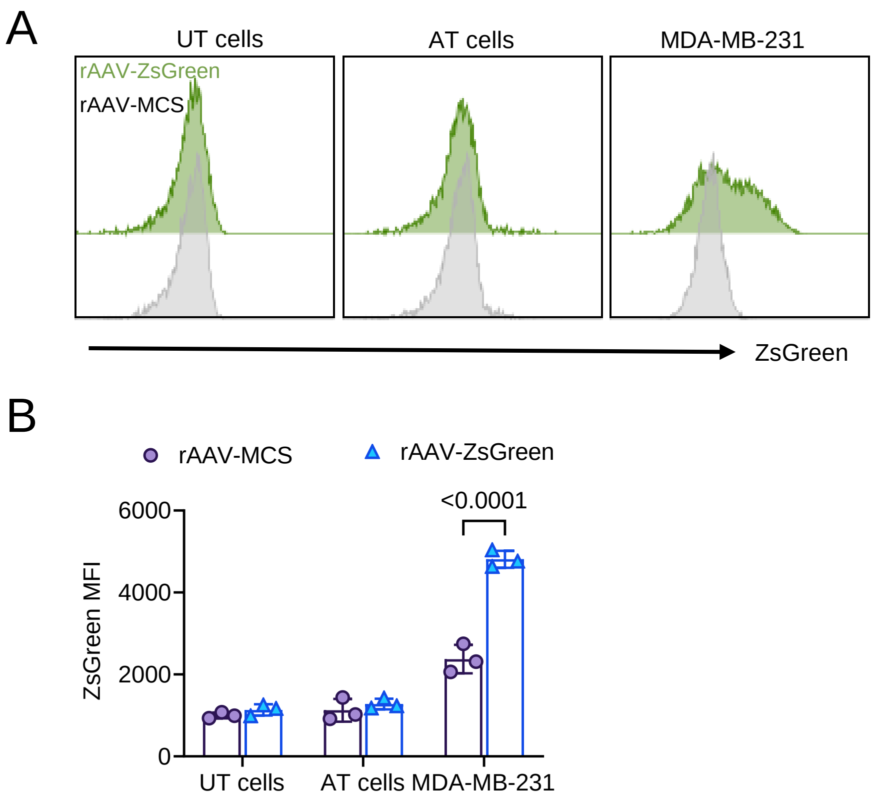


**Supplementary Figure 3. DMP-driven ZsGreen expression in T cells and MDA-MB-231 cells.** (A) Representative flow cytometry histograms of ZsGreen fluorescence in unstimulated T cells (UT), activated T cells (AT), and MDA-MB-231 cells 72 h after treatment with rAAV-MCS or rAAV-DMP-ZsGreen (rAAV-ZsGreen). (B) Quantification of ZsGreen mean fluorescence intensity (MFI) in the indicated cell populations. Data are presented as mean ± SD from three independent experiments. Statistical significance was assessed by two-way ANOVA with Šídák's multiple-comparisons test.

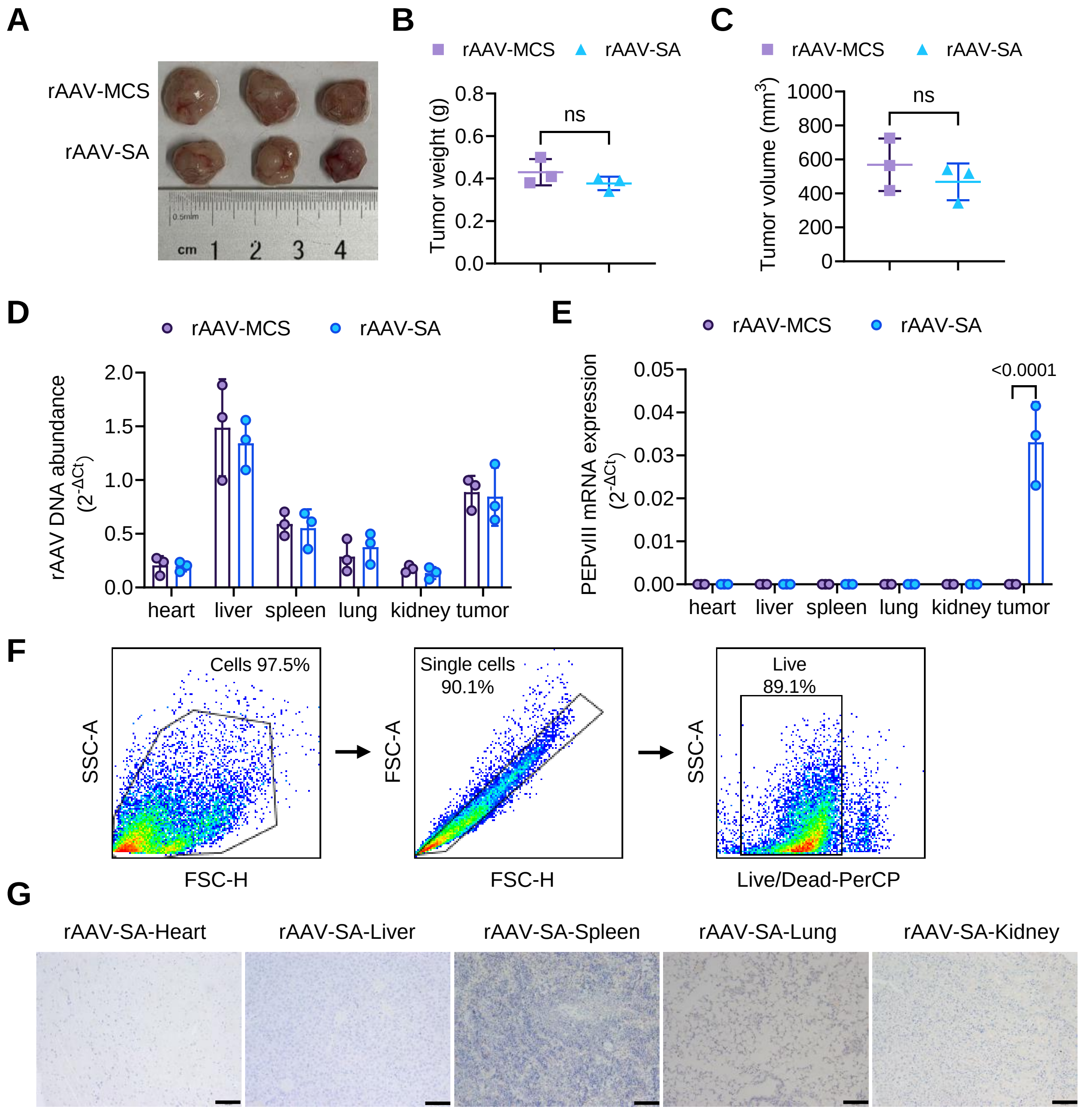


**Supplementary Figure 4. Biodistribution and tumor-preferential synthetic antigen expression following systemic rAAV-SA delivery.** MDA-MB-231 tumor-bearing NCG mice received a single intravenous administration of rAAV-MCS or rAAV-SA ($5 \times 10^{10}$ vg per mouse), and tumors and major organs were collected one week later. (A) Excised tumors from mice in each group (n = 3). (B, C) Endpoint tumor weight (B) and tumor volume (C). (D) rAAV vector DNA abundance in the heart, liver, spleen, lung, kidney, and tumor tissues determined by qPCR. (E) PEPvIII mRNA expression in the indicated tissues determined by RT-qPCR. (F) Representative flow cytometry gating strategy for analysis of surface PEPvIII expression in dissociated tumor samples. Cells were sequentially gated for cells, singlets, and viable cells before PEPvIII analysis. (G) Representative immunohistochemical staining of the Myc tag in the heart, liver, spleen, lung, and kidney from rAAV-SA-treated mice. Scale bars, 100 μm. Data are presented as mean ± SD. Statistical significance was assessed by unpaired two-tailed Student's *t*-tests (B, C) and two-way ANOVA with Šídák's multiple-comparisons test (E). ns, not significant.

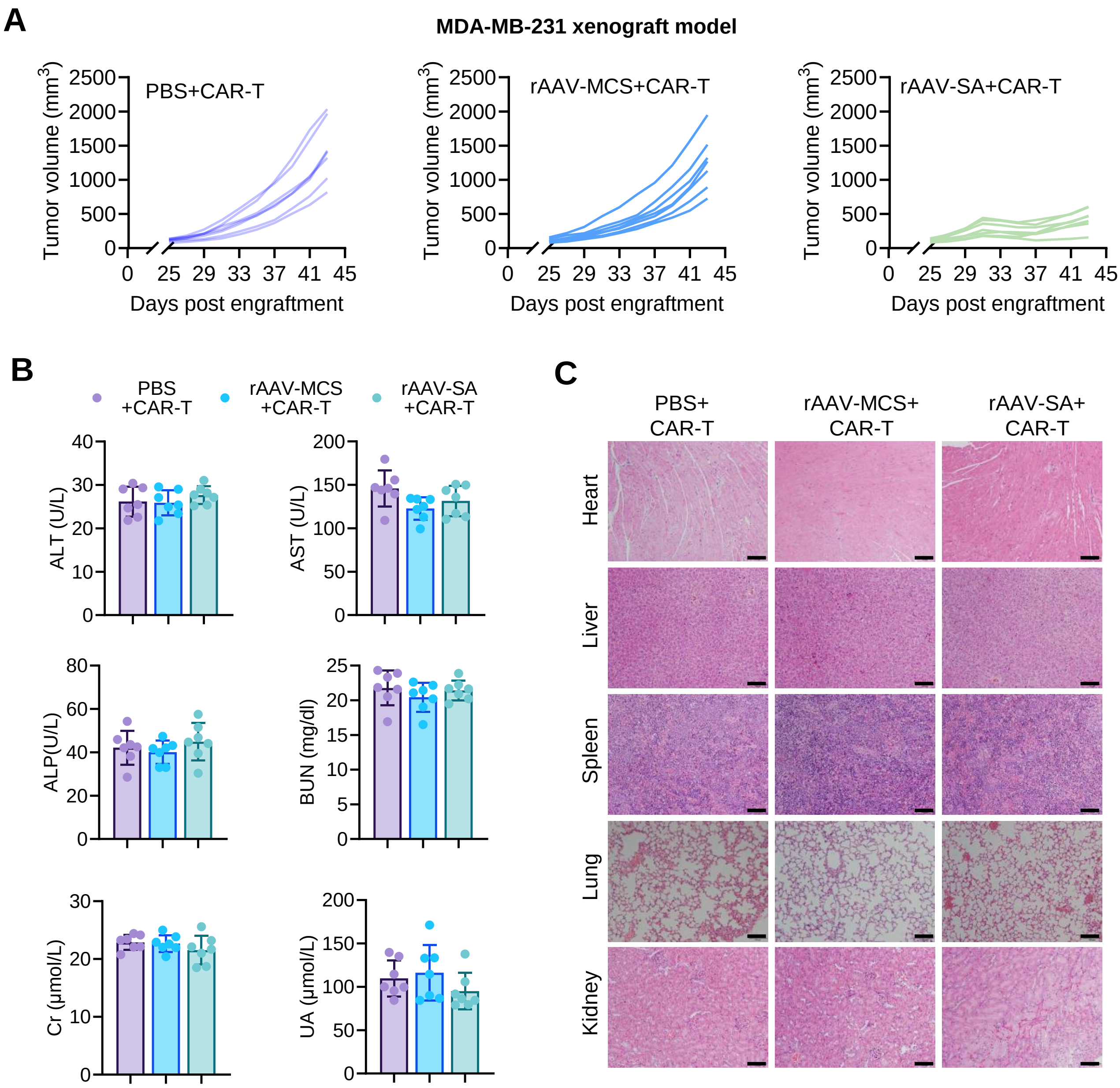


**Supplementary Figure 5. Individual tumor growth and safety evaluation in MDA-MB-231 tumor-bearing mice.** MDA-MB-231 tumor-bearing NCG mice received PBS or intravenous rAAV-MCS or rAAV-SA ($5 \times 10^{10}$ vg per mouse) on days 25, 27, and 29 after tumor engraftment, followed by intratumoral administration of CAR-T cells ($5 \times 10^{6}$ cells per mouse) on day 31. Mice were euthanized when tumor volume reached approximately 2,000 mm³, and samples were collected for endpoint analyses. (A) Individual tumor growth curves in the PBS + CAR-T, rAAV-MCS + CAR-T, and rAAV-SA + CAR-T groups. (B) Serum biochemical parameters at the endpoint, including alanine aminotransferase (ALT), aspartate aminotransferase (AST), alkaline phosphatase (ALP), blood urea nitrogen (BUN), creatinine (Cr), and uric acid (UA). (C) Representative hematoxylin and eosin (H&E) staining of the heart, liver, spleen, lung, and kidney from the indicated treatment groups. Scale bars, 100 μm. Data are presented as mean ± SD ($n$ = 7 mice per group).

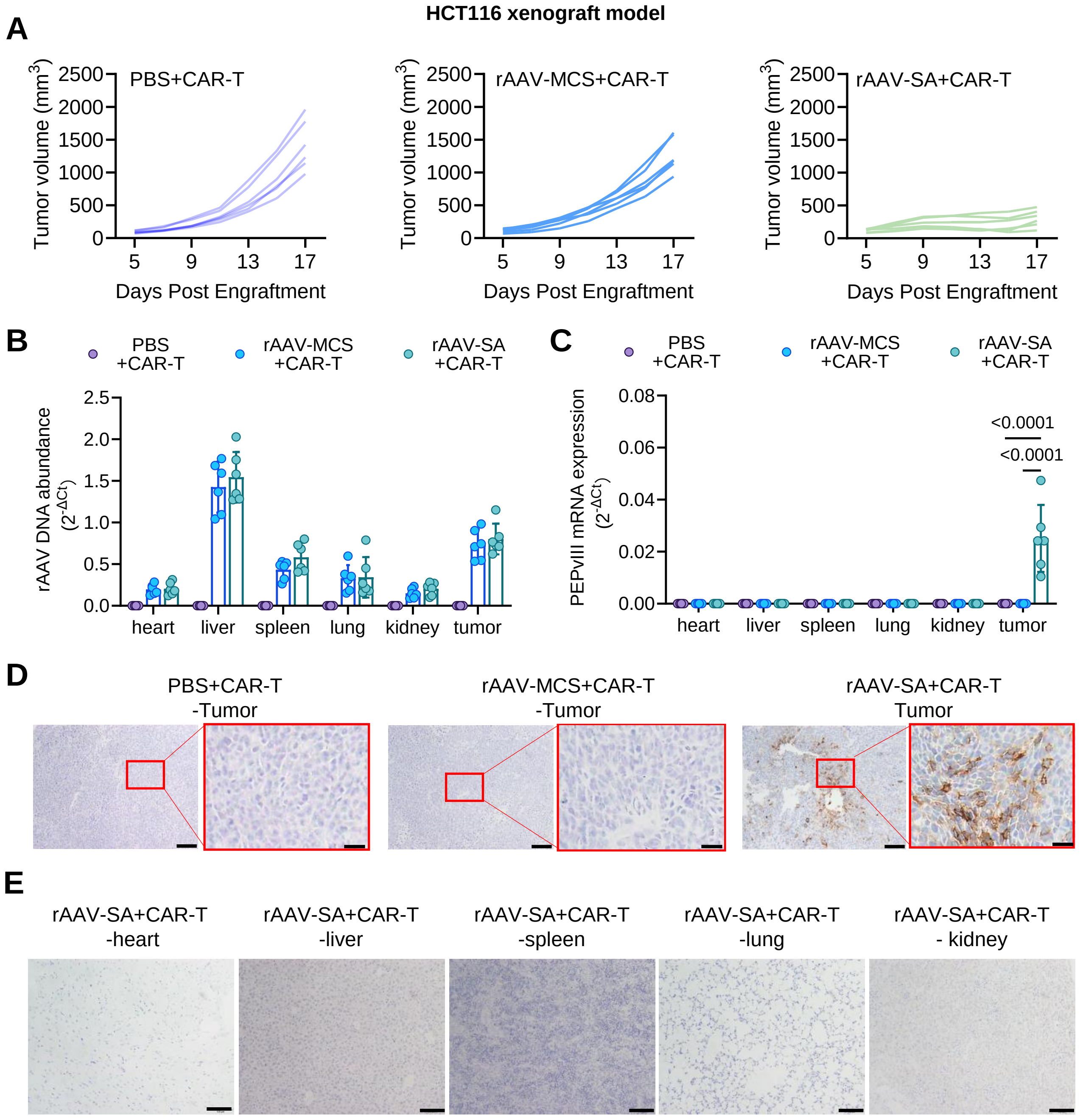


**Supplementary Figure 6. Antitumor response, rAAV biodistribution, and synthetic antigen expression in HCT116 tumor-bearing mice**. HCT116 tumor-bearing NCG mice received PBS, rAAV-MCS, or rAAV-SA ($5 \times 10^{10}$ vg/mouse) intravenously on days 5 and 7, followed by intratumoral administration of CAR-T cells ($5 \times 10^{6}$ cells/mouse) on days 9 and 13, as shown in Figure 3I. (A) Individual tumor growth curves for the indicated treatment groups. (B) rAAV vector DNA abundance in the heart, liver, spleen, lung, kidney, and tumor tissues at the experimental endpoint, determined by qPCR. (C) PEPvIII mRNA expression in the indicated tissues, determined by RT-qPCR. (D) Representative Myc-tag immunohistochemistry images of tumor tissues from the indicated treatment groups. Scale bars, 100 μm for overview images and 25 μm for magnified images. (E) Representative Myc-tag immunohistochemistry images of the heart, liver, spleen, lung, and kidney from rAAV-SA + CAR-T-treated mice. Scale bars, 100 μm. Data are presented as mean ± SD (n = 6 mice per group). Statistical significance was assessed by two-way ANOVA with Šídák's multiple-comparisons test (C).

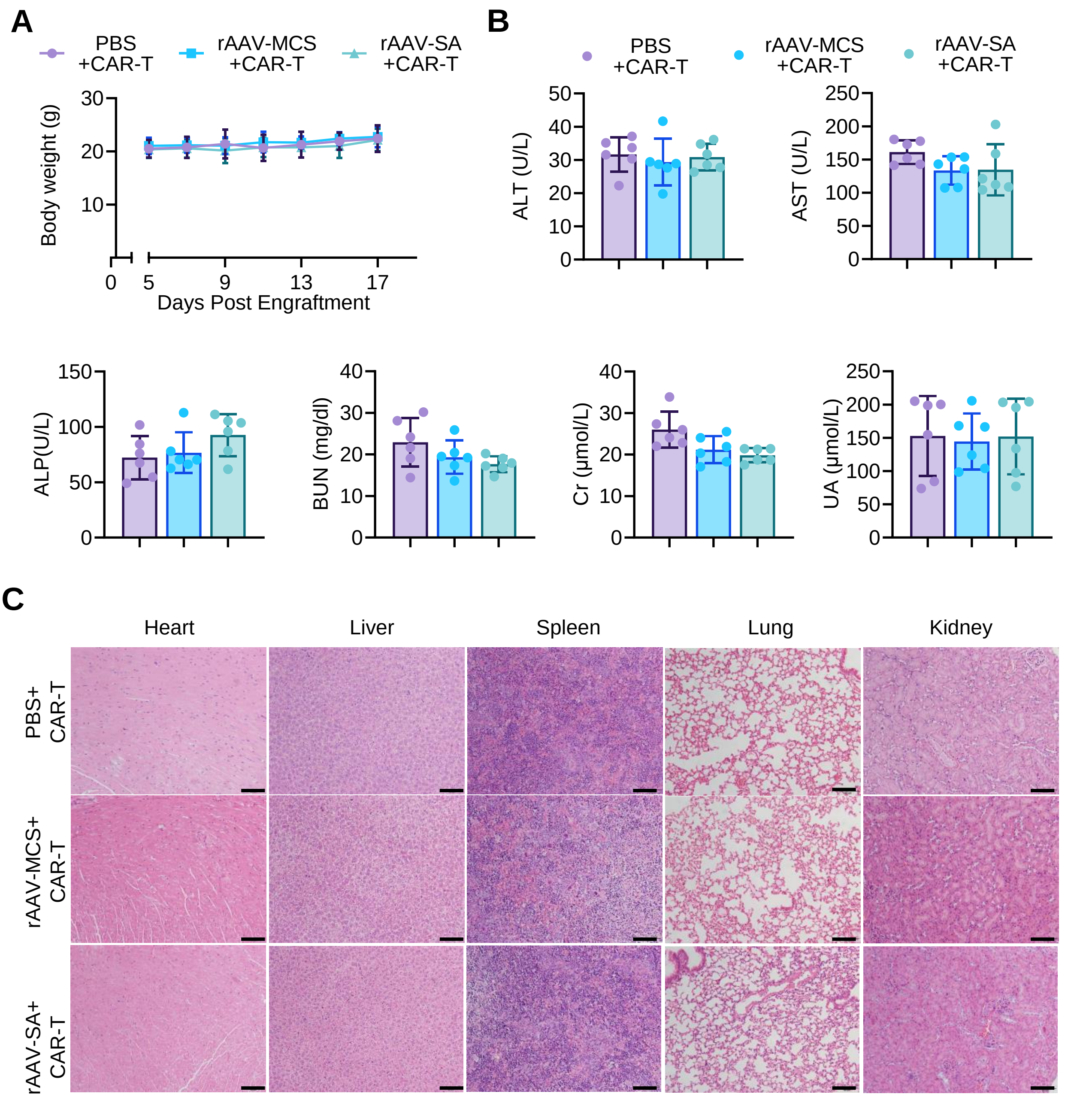


**Supplementary Figure 7. Safety evaluation of rAAV-SA combined with CAR-T-cell treatment in HCT116 tumor-bearing mice.** HCT116 tumor-bearing NCG mice were treated according to the schedule shown in Figure 3I. **(A)** Body-weight changes during treatment. **(B)** Serum biochemical parameters measured at the experimental endpoint, including ALT, AST, ALP, BUN, Cr, and UA. **(C)** Representative H&E staining of the heart, liver, spleen, lung, and kidney from the indicated treatment groups. Scale bars, 100 μm. Data are presented as mean ± SD ($n$ = 6 mice per group).

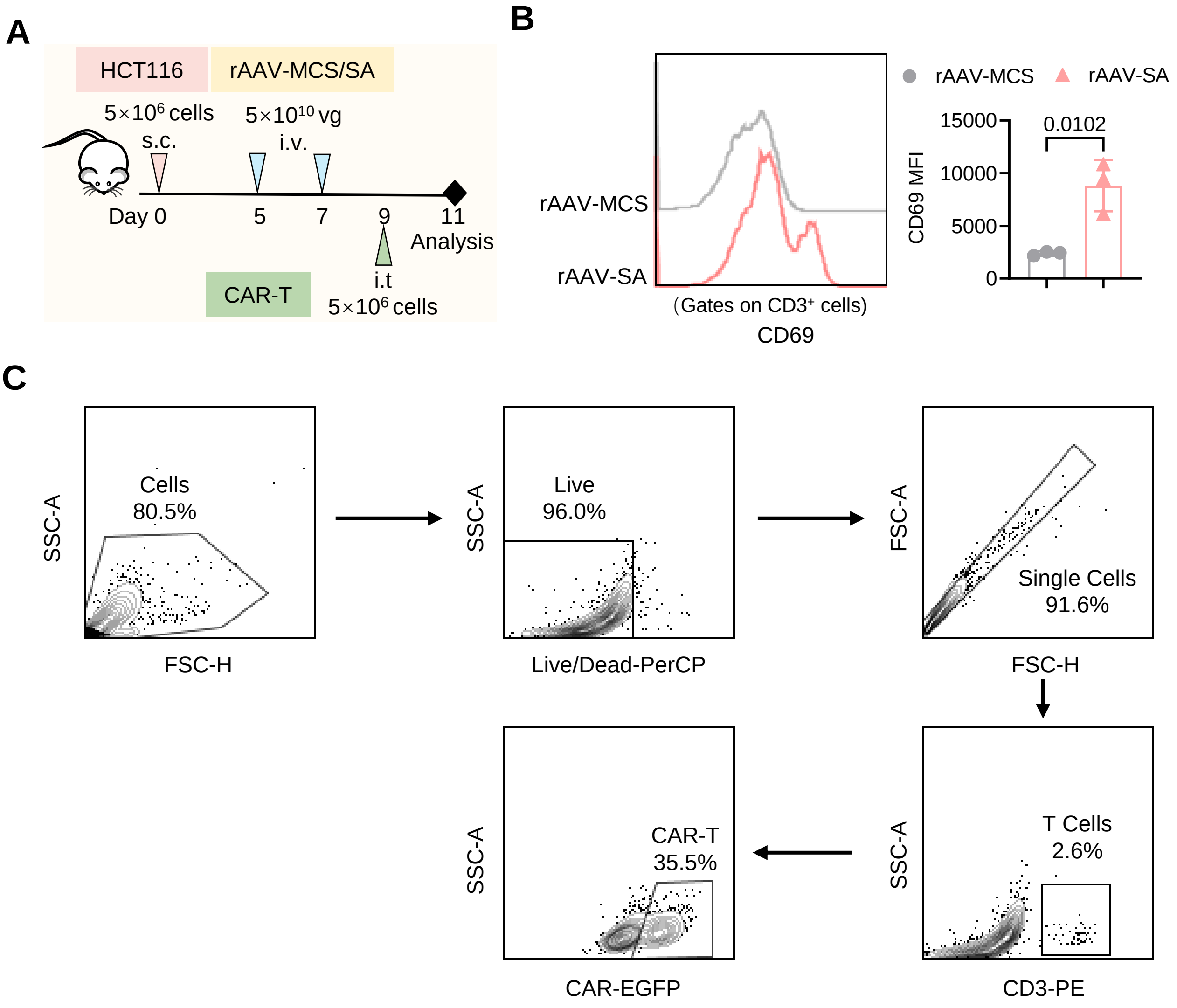


**Supplementary Figure 8. Intratumoral T-cell activation and flow cytometry gating strategy in HCT116 tumor-bearing mice. (A)** Experimental scheme for short-term analysis of intratumoral T-cell activation. HCT116 tumor-bearing NCG mice received intravenous rAAV-MCS or rAAV-SA on days 5 and 7, followed by intratumoral CAR-T-cell administration on day 9. Tumors were analyzed two days later. **(B)** Representative flow cytometry histograms and quantification of CD69 mean fluorescence intensity (MFI) in intratumoral $CD3^+$ T cells. **(C)** Representative flow cytometry gating strategy for identification of intratumoral $CAR^+$ T cells. Cells were sequentially gated for cells, viable cells, singlets, $CD3^+$ T cells, and $CAR^+$ T cells. Data in (B) are presented as mean ± SD (*n* = 3 mice per group). Statistical significance was assessed using an unpaired two-tailed Student's *t*-test.

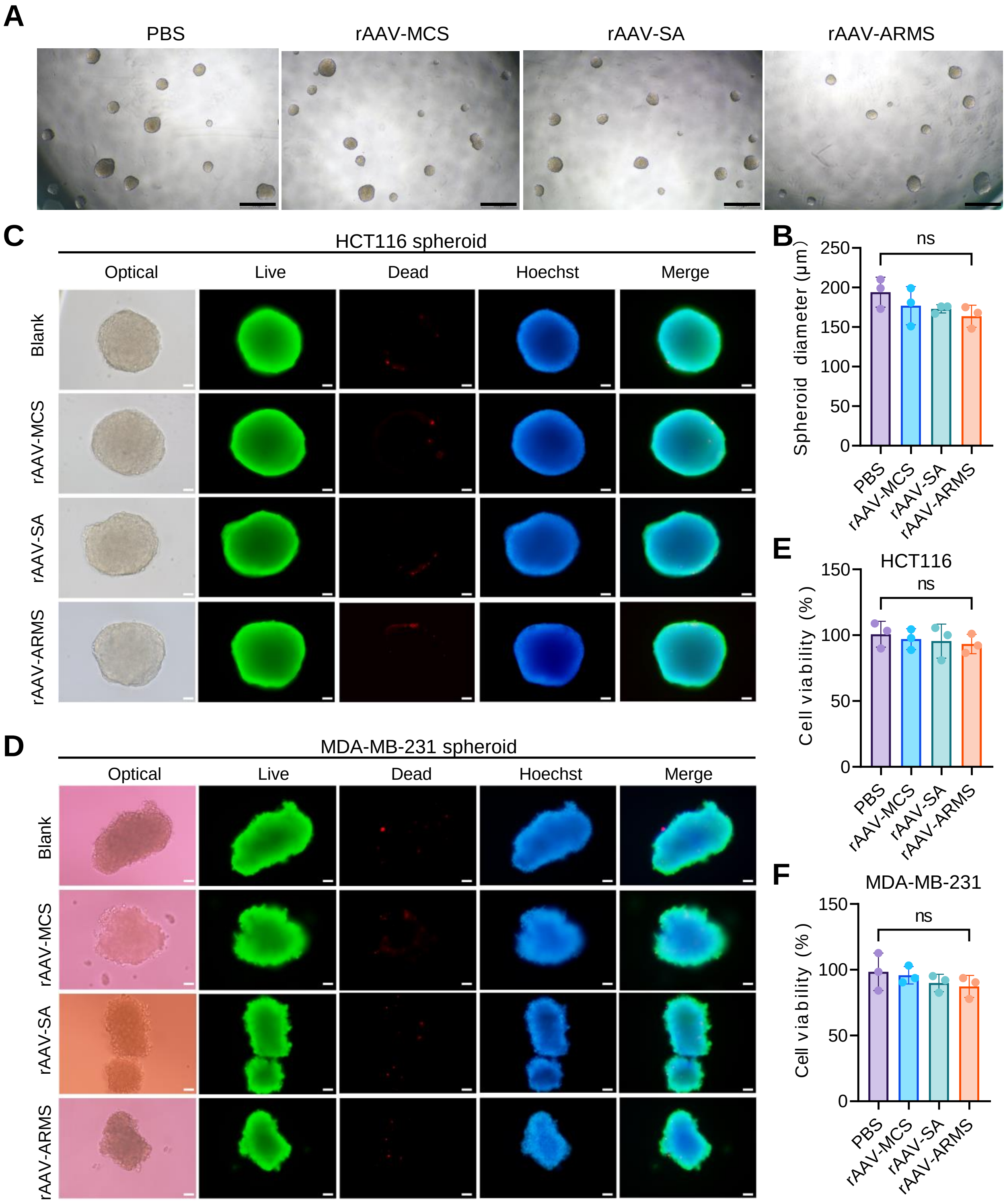


**Supplementary Figure 9. rAAV treatment alone does not affect tumor spheroid growth or viability. (A)** Representative bright-field images of HCT116 spheroids 72 h after treatment with PBS, rAAV-MCS, rAAV-SA, or rAAV-ARMS. Scale bars, 500 µm. **(B)** Quantification of HCT116 spheroid mean diameter. **(C, D)** Representative fluorescence images of HCT116 (C) and MDA-MB-231 (D) spheroids stained with Calcein AM, PI, and Hoechst after the indicated rAAV treatments. Scale bars, 50 µm. **(E, F)** Luciferase-based viability analysis of HCT116-Luc (E) and MDA-MB-231-Luc (F) spheroids following the indicated rAAV treatments. Data are presented as mean ± SD from three independent experiments. Statistical significance was assessed by one-way ANOVA with Tukey's multiple-comparisons test.

.

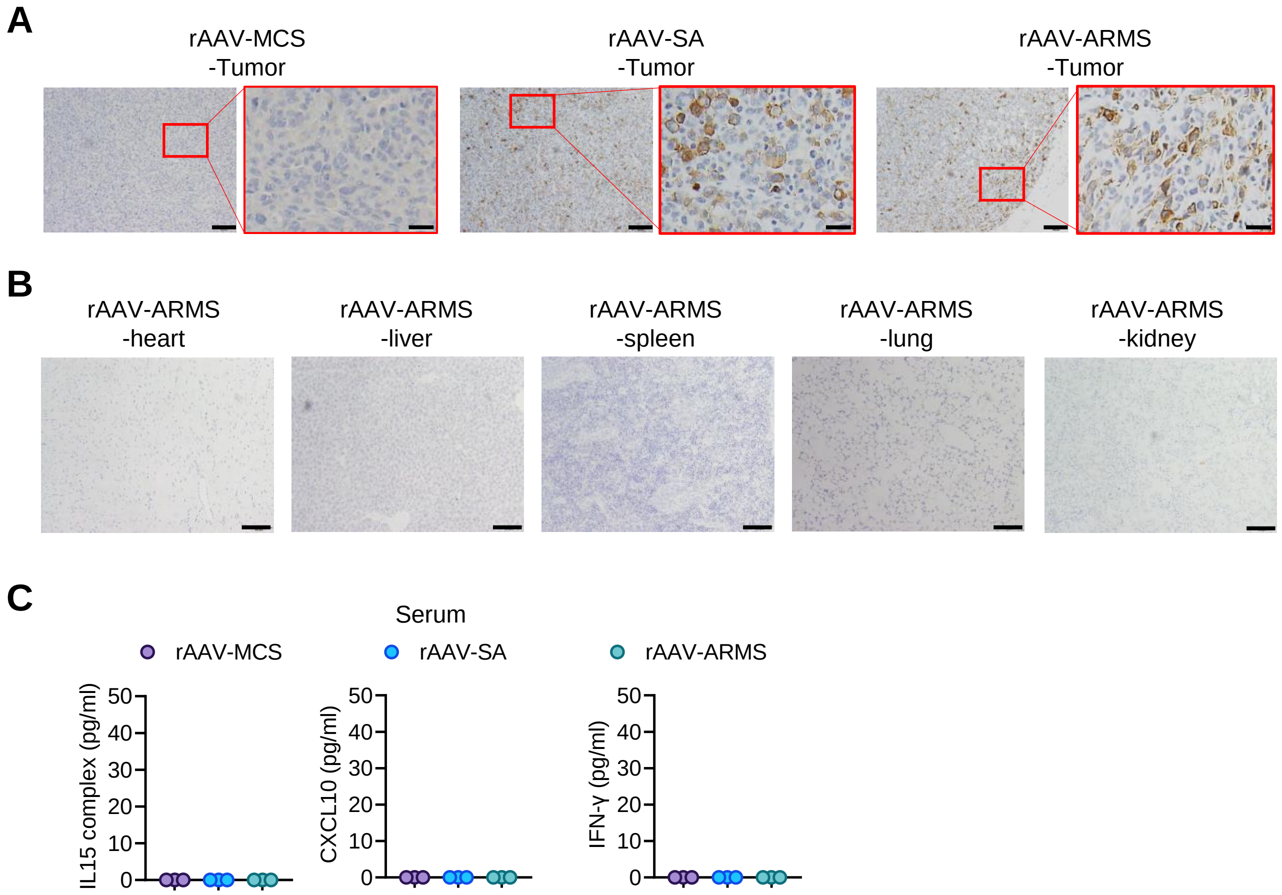


**Supplementary Figure 10**. **Tumor transgene expression and systemic exposure of rAAV-ARMS payloads in MDA-MB-231 tumor-bearing mice.** MDA-MB-231 tumor-bearing NCG mice received a single intravenous injection of rAAV-MCS, rAAV-SA, or rAAV-ARMS ($1 \times 10^{11}$ vg/mouse) on day 22, and samples were collected one week later. (A) Representative Myc immunohistochemistry images of tumor tissues from the indicated treatment groups. Scale bars, 100 μm for overview images and 25 μm for magnified images. (B) Representative Myc immunohistochemistry images of the heart, liver, spleen, lung, and kidney from rAAV-ARMS-treated mice. Scale bars, 100 μm. (C) Serum concentrations of IL-15 complex, CXCL10, and IFN-γ in the indicated treatment groups. All three factors were below the assay detection limit under the conditions tested. Data are presented as mean ± SD (n = 3 mice per group).

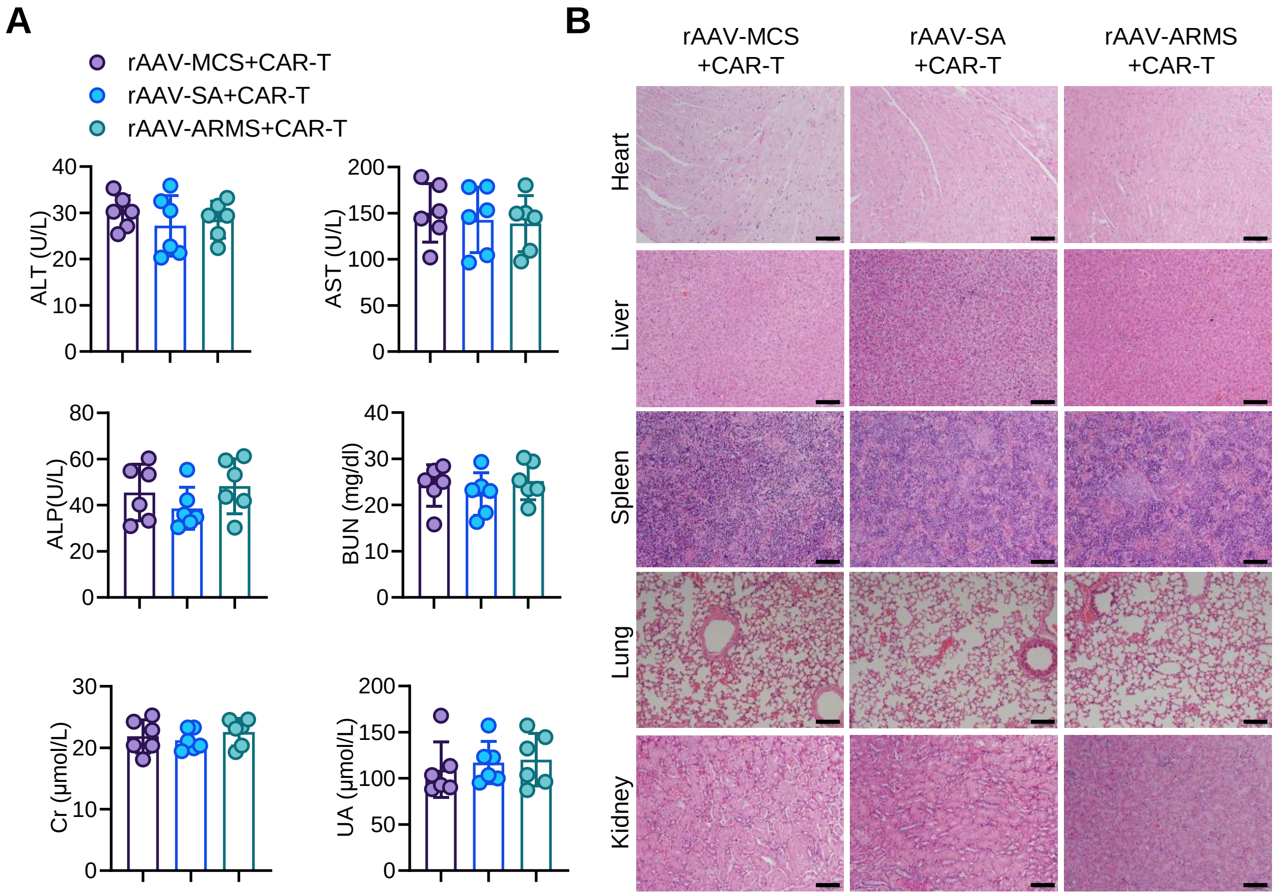


**Supplementary Figure 11. Systemic safety evaluation of rAAV-ARMS combined with CAR-T-cell therapy in MDA-MB-231 tumor-bearing mice.** MDA-MB-231 tumor-bearing NCG mice received a single intravenous injection of rAAV-MCS, rAAV-SA, or rAAV-ARMS ($1 \times 10^{11}$ vg per mouse) on day 22, followed by intravenous CAR-T-cell infusions ($5 \times 10^{6}$ cells per mouse) on days 27 and 34. Samples were collected at the experimental endpoint. **(A)** Serum biochemical parameters, including alanine aminotransferase (ALT), aspartate aminotransferase (AST), alkaline phosphatase (ALP), blood urea nitrogen (BUN), creatinine (Cr), and uric acid (UA). **(B)** Representative hematoxylin and eosin (H&E) staining of the heart, liver, spleen, lung, and kidney from the indicated treatment groups. Scale bars, 100 μm. Data are presented as mean ± SD (*n* = 6 mice per group). Statistical significance was assessed using one-way ANOVA with multiple-comparisons testing.
